# A generalisable method for the purification and biophysical characterisation of bacterial membrane receptors

**DOI:** 10.64898/2026.08.12.744447

**Authors:** Federico Bosetto, Maria Zacharopoulou, Mark Bycroft, Monika Kish, Giovanna Zinzalla, Pamela J. E. Rowling, Stephen H. McLaughlin, Jonathan J. Phillips, Laura S. Itzhaki, Ioanna Mela

## Abstract

Membrane-embedded bacterial receptors are challenging to express and purify in soluble form, yet their isolated domains are essential tools for structural and ligand-discovery studies. *Pseudomonas aeruginosa* relies on the TonB-dependent heme receptor HasR for iron acquisition, a process central to its pathogenicity. Here, we report a robust strategy for the recombinant expression, purification, and biophysical characterisation of the two soluble HasR domains directly involved in heme uptake: the N-terminal plug and the Secretin/TonB short N-terminal domain. Each domain was expressed individually in *E. coli* and purified to homogeneity, adopting well-folded conformations as confirmed by circular dichroism, NMR spectroscopy, and mass spectrometry. We then engineered a fusion construct containing both domains and systematically evaluated multiple solubilisation tags. A GST-His dual-affinity strategy enabled efficient purification of the construct, whereas His-tag alone resulted in insoluble protein and HLT-tag fusions suffered from non-specific proteolysis. Biophysical analyses revealed that the Secretin/TonB short N-terminal domain remains stably folded within the fusion construct, while the N-terminal plug domain becomes partially disordered, a finding further supported by hydrogen/deuterium exchange mass spectrometry. Together, these results establish a generalizable workflow for producing soluble receptor domains from membrane proteins and provide validated HasR constructs suitable for downstream ligand-screening applications, including aptamer and nanobody discovery.

## Introduction

*Pseudomonas aeruginosa* is a Gram-negative bacterium, known for its remarkable capability to survive in diverse environments, as it can thrive on minimal nutrients and can tolerate a broad range of environmental conditions^1,2^. This adaptability allows it to persist in both community settings and, notably, in the challenging environment of healthcare facilities, where it appears as an opportunistic pathogen causing infection and mortality in cystic fibrosis patients and immunocompromised individuals^1,2,3^. *P. aeruginosa* is resistant to several classes of antibiotics, including aminoglycosides, quinolones and β-lactams, due to a combination of different factors including intrinsic, acquired and adaptive resistance mechanisms^2,3,4^. Antibiotic resistance shown by *P. aeruginosa* is due to intrinsic and acquired traits, such as low permeability of the outer membrane, efflux pumps that actively transport antibiotics out of the cell, and the production of β-lactamases and aminoglycoside-modifying enzymes that impair antibiotic activity. Furthermore, *P. aeruginosa* can form biofilms, which act as a physical barrier to antibiotic penetration into bacterial cells^2,3,4^. *P. aeruginosa* belongs to the group of ESKAPE pathogens, represented by the six most multidrug-resistant bacteria and for which the development of new treatments is necessary, including: *Enterococcus faecium, Staphylococcus aureus, Klebsiella pneumoniae, Acinetobacter baumannii, Pseudomonas aeruginosa* and *Enterobacter* spp^5^.

Iron is essential for bacterial growth^,7^, and plays a significant role in various cellular functions, such as energy production, DNA replication, electron transport, and virulence^8^. In vertebrates, iron is mainly coordinated within a porphyrin ring structure named heme^9,10,11^ which is responsible of the transport and storage of oxygen in proteins such as haemoglobin and myoglobin^9,11^. Sequestering heme within proteins is a well-documented strategy adopted by vertebrates to prevent bacterial infections, however, many pathogens have developed mechanisms to utilize heme as a source of iron^9,11^. In *P. aeruginosa*, three main heme uptake systems are known: the heme assimilation system (*Has*)^6,7,10,12^, the *Pseudomonas* heme uptake (*Phu*)^6,7,12^ and the TonB-dependent *Hxu* system^13^. The Has pathway is unique in its reliance on the secreted hemophore HasA, which captures extracellular heme and delivers it to the outer-membrane receptor HasR (Figure 1.A)^6,7,12,14^, whereas Phu ^6,7,14^ and Hxu ^13^ receptors bind free heme directly.

**Figure 1.**
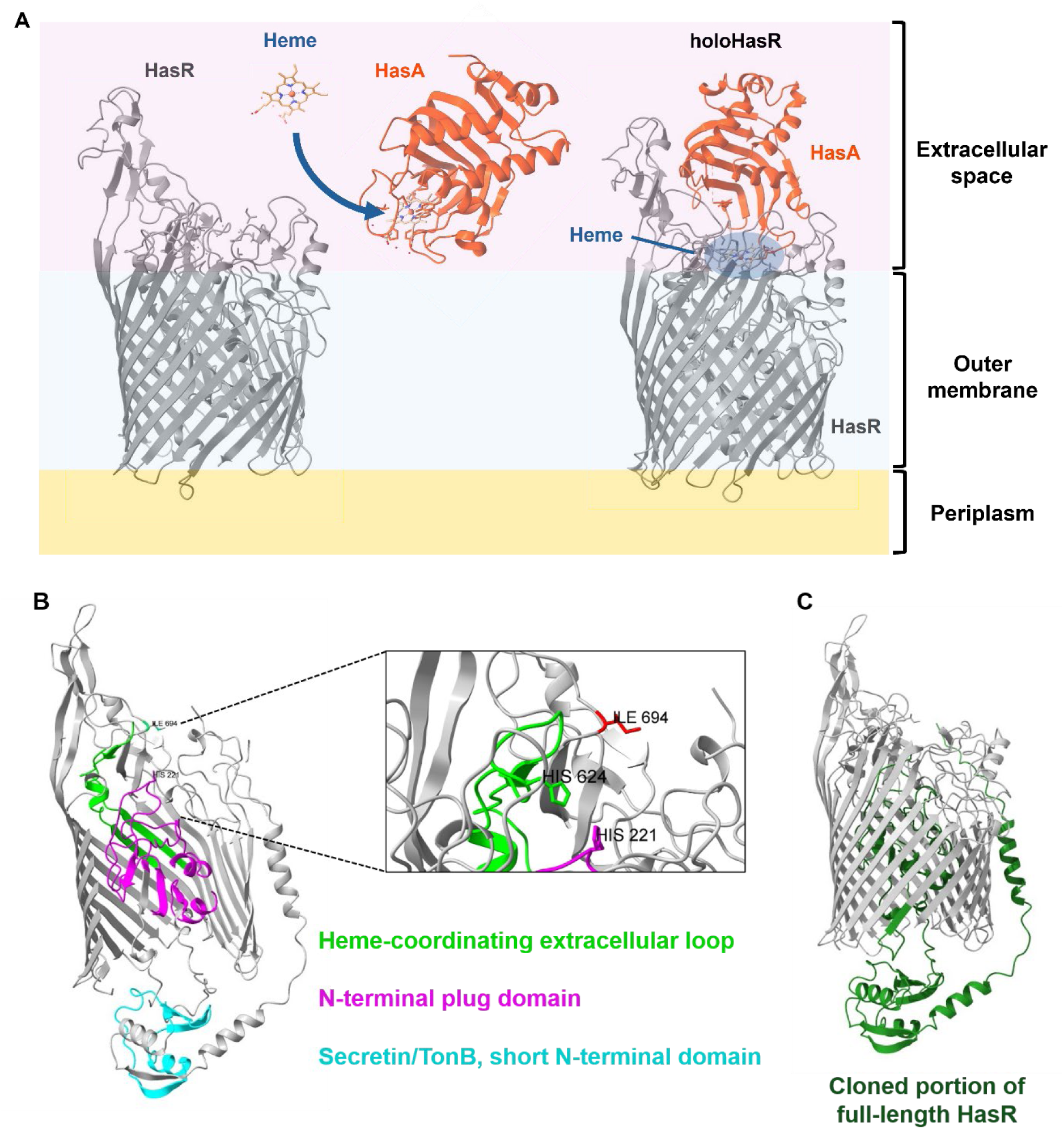
**(A)** Schematic representation of the heme assimilation system in *Pseudomonas aeruginosa*, constituted by the receptor HasR (in grey) and the hemophore HasA (in orange) which binds the heme (blue arrow), and together form the holoHasR. The image was adapted from PDB file 3CSL. **(B)** Illustration of the three HasR domains: Secretin/TonB short N-terminal (in cyan), N-terminal plug (in purple), and heme-coordinating extracellular loop (in light green); where the last two are directly involved in the heme coordination binding thanks to His-221 and His-624 residues. The beta barrel structure is partially concealed to allow for visualisation of the domains. **(C)** Highlight of the cloned portion of HasR including both N-terminal plug and Secretin/TonB short N-terminal domains (in dark green). The image was generated with AlphaFold2.

HasR, (Figure 1.A), is a TonB-dependent transporter composed of three domains: a short N-terminal Secretin/TonB domain, an N-terminal plug domain, and an extracellular heme-coordinating loop (Figure 1.B); The plug and extracellular loop contain key residues—His-221, His-624, and Ile-694—that mediate heme transfer from holo-HasA, regulate transport, and prevent heme back-transfer to HasA ^7^. The importance of iron acquisition is central to bacterial survival, the soluble domains of HasR represent attractive targets for therapeutic ligand discovery.

Although full-length membrane receptors are often difficult to express in soluble form due to hydrophobic transmembrane regions^15^, their isolated extracellular and periplasmic domains can frequently be produced recombinantly and purified under native conditions^16, 17^. Such soluble domains are valuable tools for structural studies and for screening ligands—including aptamers or nanobodies—that may disrupt receptor function. However, expressing isolated domains of membrane proteins presents challenges: domain boundaries must be carefully selected ^18,19^, solubility and folding can be compromised outside the native membrane environment, and purification often requires tailored strategies or solubilisation tags.

A carefully designed purification strategy is critical for ensuring yield, purity, and functionality. Affinity tags such as His, Glutathione S-Transferase (GST), Small Ubiquitin-like Modifier (SUMO), or Maltose Binding Protein (MBP) can enhance solubility and folding, especially when fused to the N-terminus of the protein of interest ^20,21^. Chromatographic techniques—including ion exchange (IEX), hydrophobic interaction (HIC), and size-exclusion chromatography (SEC)—are frequently required to achieve high purity. SEC is especially valuable not only for removing impurities and aggregates but also for analysing the size and oligomeric state of the protein of interest (monomer/oligomers)^22^. Expressing and purifying only the soluble domain of the protein of interes allows one to avoid the use of detergents entirely^23^ while preserving structural integrity. However, if residual hydrophobic patches remain, mild detergents are sometimes necessary. Strategies to increase overall protein yield include lower induction temperatures, reduced IPTG levels, and initiating induction during late log phase^24,25^, matching codon usage to the host organism, or co-expressing chaperones^26^. Once purified, biophysical methods such as circular dichroism, nanoDSF, NMR, and ligand-binding assays are essential to confirm correct folding and functionality.

Here we report a rapid and robust protocol for the expression and purification of the soluble domains of HasR: the N-terminal plug, the Secretin/TonB short N-terminal domain, and a fusion construct containing both (Figure 1.B and C). The two domains were first expressed and characterised individually, and subsequently as a single contiguous construct, reflecting their proximity within the amino-acidic sequence of HasR (Supplementary Figure S1.A and B). The heme-coordinating extracellular loop, also involved in heme binding, was not included as it is separated from these domains by the membrane-embedded β-barrel and is not accessible to targeting molecules. We evaluate multiple fusion-tag strategies, optimise expression and solubility, and establish a purification workflow combining affinity chromatography, size-exclusion chromatography, and ion-exchange polishing. Finally, we characterise the folding and stability of the purified domains using CD, nanoDSF, NMR, and HDX-MS. Our work provides validated HasR constructs suitable for downstream ligand-discovery applications and offers a generalizable strategy for producing soluble domains of membrane-associated bacterial receptors.

## Results

### Recombinant individual HasR domains can be purified and are soluble

To determine whether the soluble regions of HasR could be produced recombinantly, we first expressed the N-terminal plug and Secretin/TonB short N-terminal domains individually (Figure 1.B and C and Supplementary Figure S1.A and B). Both proteins were soluble in *E. coli* lysates, although the Secretin/TonB domain expressed at substantially higher levels.

The N-terminal plug domain was purified through a three-step purification process due to the presence of high amount of host-derived contaminants. The first step was performed via nickel affinity chromatography resulting in a well-defined protein band localised between 23 kDa and 18 kDa, but with protein contaminants at lower and higher molecular weight (Figure 2.A and B). The second purification step of anion-exchange chromatography (ANEX) removed a portion of host-derived contaminants (Figure 2.C and D) and the third and final purification by SEC produced a single major peak corresponding to the plug domain (Figure 2.E and F). ESI mass spectrometry confirmed a dominant species at 16.9 kDa, consistent with the predicted mass of 15.8 kDa (Supplementary Figure S2.A). The total yield of the N-terminal plug domain resulted in ∼0.075 mg/L (0.45 mg from 6 liters).

**Figure 2.**
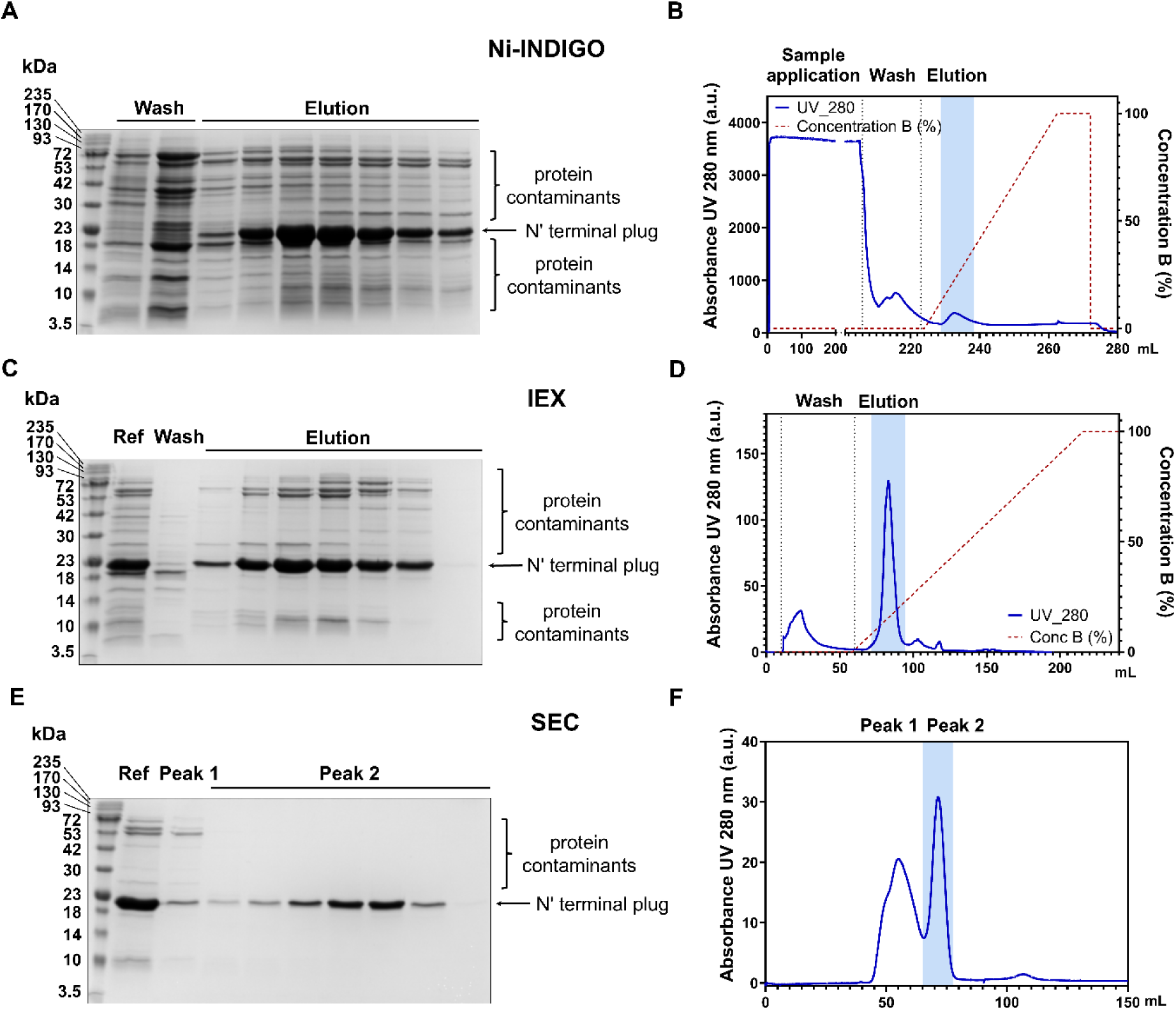
N-terminal plug domain purification. The protein is initially purified by affinity chromatography using Ni-INDIGO column as reported by **(A)** the SDS-PAGE and **(B)** the chromatogram from the AKTA. N-terminal plug protein band is denoted by arrow between 23 kDa and 18 kDa, protein contaminants are present both at higher and lower molecular weight. The chromatogram reports Sample application, Wash and Elution phases; elution peak is highlighted by blue shadow. Second purification step was carried out via IEX using a MonoQ column, again **(C)** SDS-PAGE and **(D)** the chromatogram from the AKTA are shown. N-terminal plug protein band is denoted by arrow between 23 kDa and 18 kDa, some protein contaminants are still present both at higher and lower molecular weight. The chromatogram reports Wash and Elution phases; elution peak from N-terminal plug is highlighted by blue shadow. The final step was performed by SEC using a Superdex 75 10/300 GL column, which separated the protein of interest from residual contaminants, as seen in **(E)** the SDS-PAGE and **(F)** AKTA chromatogram. N-terminal plug protein band is denoted by arrow between 23 kDa and 18 kDa (Peak 2), while a few protein contaminants are still present at higher molecular weight (Peak 1). The chromatogram reports Peak 1 and Peak 2; elution peak from N-terminal plug is highlighted by blue shadow (Peak 2).

The Secretin/TonB short N-terminal domain was initially purified by nickel affinity chromatography, yielding a clear, defined protein band at approximately 10 kDa (Figure 3.A and B). SEC removed remaining impurities, yielding a homogeneous protein. (Figure 3.C and D). The purity and correct molecular weight of the purified protein of 13.1 kDa was confirmed via ESI mass spectrometry (Supplementary Figure S2.B). The total yield of the Secretin/TonB short N-terminal domain resulted in ∼12.3 mg/L (12.3 mg from 1 liter).

**Figure 3.**
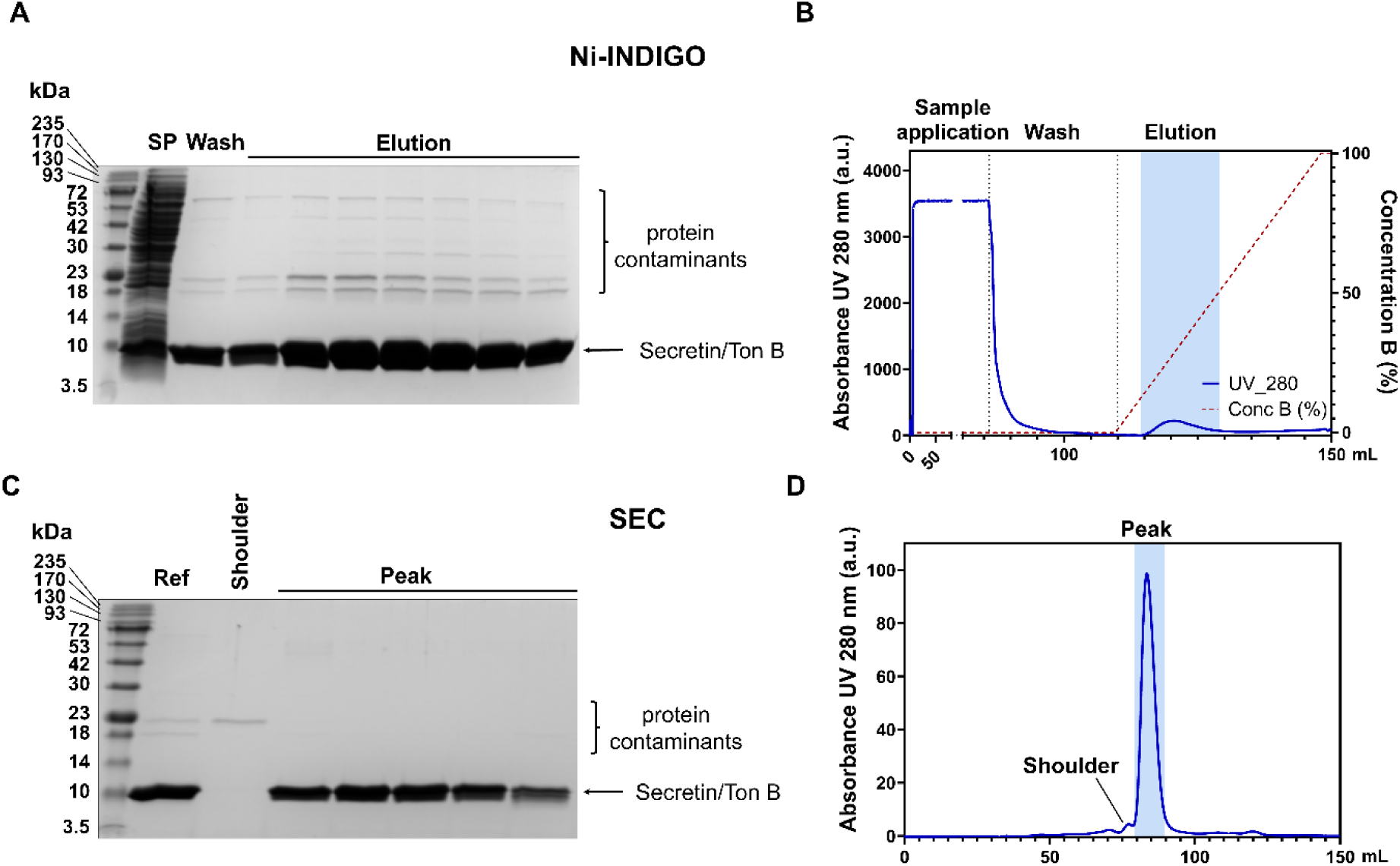
Secretin/TonB short N-terminal domain purification. The protein is initially purified by affinity chromatography using Ni-INDIGO column as reported by **(A)** the SDS-PAGE and **(B)** the chromatogram from the AKTA. Secretin/TonB short N-terminal protein band is denoted by arrow around 10 kDa, protein contaminants are present at higher molecular weight. The chromatogram reports Sample application, Wash and Elution phases; elution peak is highlighted by blue shadow. Second purification step was carried out via SEC using a Superdex 75 10/300 GL column (Cytiva, UK) to isolate the protein of interest, again **(C)** SDS-PAGE and **(D)** the chromatogram from the AKTA are shown. Secretin/TonB short N-terminal protein band is denoted by arrow around 10 kDa (Peak), and complete removal of protein contaminants is visible (Shoulder). The chromatogram reports Peak and Shoulder; elution peak from Secretin/TonB short N-terminal protein is highlighted by blue shadow (Peak). [SP= soluble protein fraction].

### Individual HasR domains adopt well-folded conformations

CD analysis revealed distinct secondary-structure profiles for the two domains. The N-terminal plug domain exhibited a mixed architecture dominated by antiparallel β-strands (52.4%), with 20.5% distorted α-helices and 27.1% right-twisted antiparallel β-strands (Figure 4.A and C), while the Secretin/TonB short N-terminal domain was almost entirely α-helical (93.5%) (Figure 4.B and D). These observations were further supported by 1D NMR spectra. The 1D NMR spectrum of the N-terminal plug is well dispersed, characteristic of a fully folded protein (e.g. peaks with low chemical shift values, between -0.5 and 0.5 ppm are resolved) (Figure 5.A). Interestingly, the resolved NH signals that are present only when the plug domain is folded are typical of β-sheets. Similarly, the 1D NMR spectrum of the Secretin/TonB short N-terminal domain showed well-dispersed spectra, characteristic of a well-folded structure (Figure 5.B). These observations were further substantiated by a 2D NOESY experiment, where widely dispersive peaks indicated long-range, through-space contacts that are only present in a folded structure (Supplementary Figure S.3).

**Figure 4.**
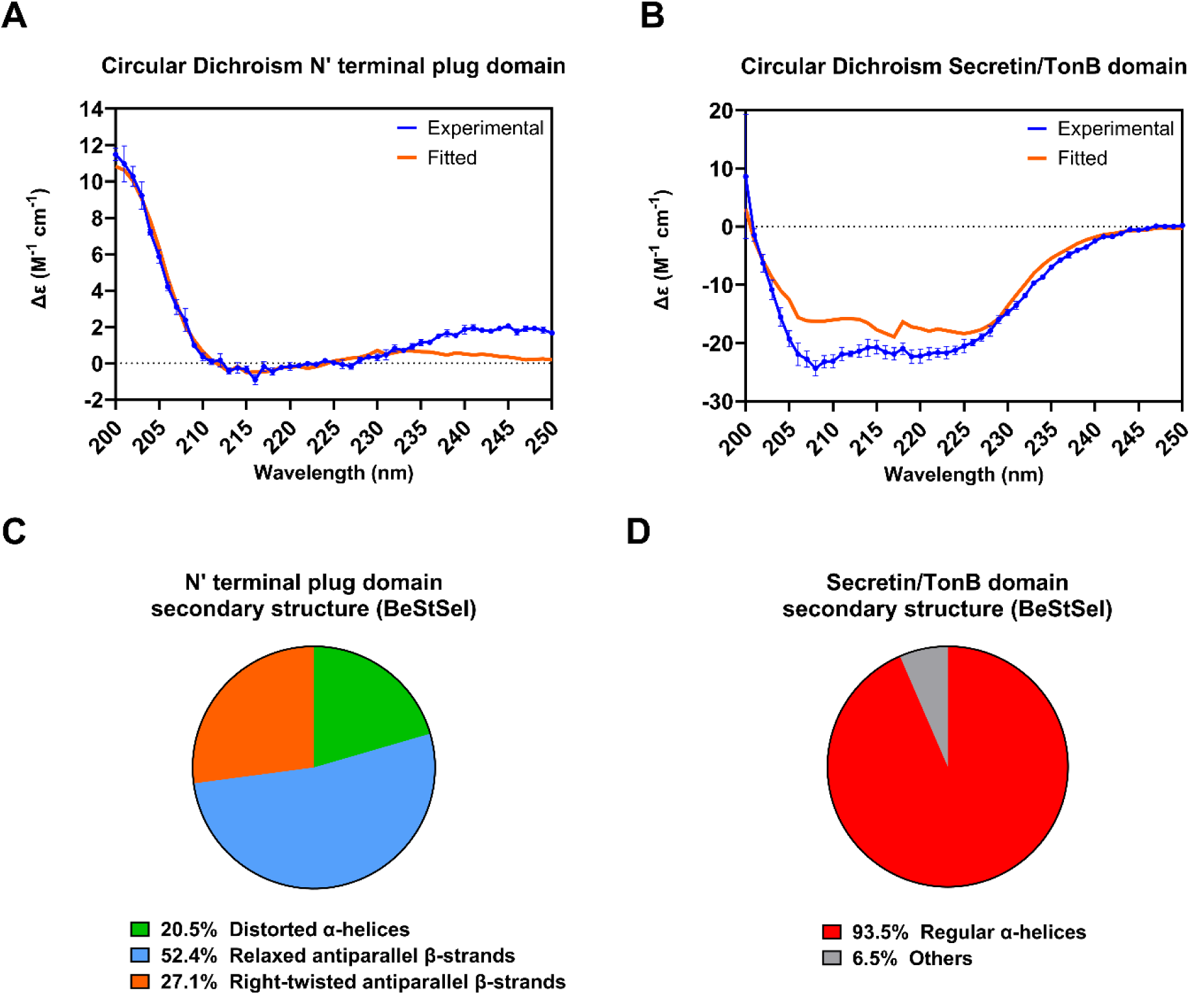
Circular Dichroism spectra of the N-terminal plug **(A)** and Secretin/TonB short N-terminal domains **(B)** were measured in triplicate at 25°C between 200 and 250 nm (in blue), the fitted curves from BeStStel are also reported (in orange). **(C)** Secondary structure prediction of the N-terminal plug domain by BeStSel server using the CD values, indicating a composition of 20.5% of distorted α-helices (dark green), 52.4% of relaxed antiparallel β-strands (light blue) and 27.1% of right-twisted antiparallel β-strands (orange). **(D)** Secondary structure prediction of the Secretin/TonB short N-terminal domain by BeStSel server using the CD values, indicating a composition of 93.5% of regular α-helices (red) and 6.5% of others (disordered, grey).

**Figure 5.**
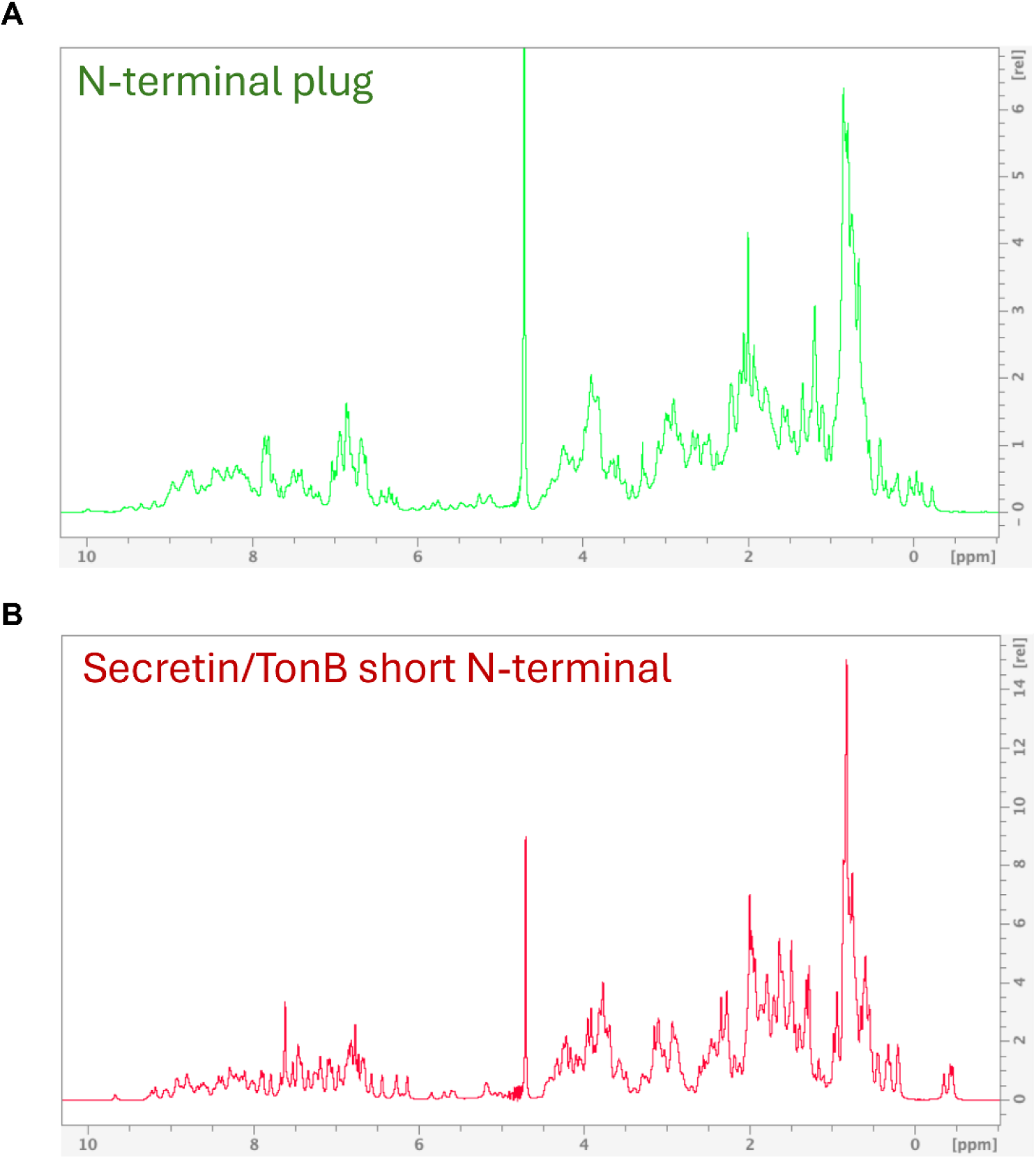
1D NMR spectra of individual N-terminal plug and Secretin/TonB short N-terminal domains. **(A)** 1D NMR spectrum of individual N-terminal plug domain. **(B)** 1D NMR spectrum of individual Secretin/TonB short N-terminal domain.

Together, these data demonstrate that both HasR domains are independently soluble and adopt well-defined tertiary structures.

### Recombinant HasR construct requires a solubilisation tag fusion

We next engineered a construct containing the N-terminal plug and the Secretin/TonB short N-terminal domains in their native sequence order (Figure 1.B and C and Supplementary Figure S1.A and B). We evaluated three fusion-tag strategies to optimise expression and solubility. The HasR dual-domain construct was fused to either a simple N-terminal 6-His tag, an N-terminal Histidine–Lipoyl (HLT) solubilisation tag, or an N-terminal Glutathione S-Transferase (GST) tag combined with a C-terminal 6-His tag (Supplementary Figure S4.A–C). Each construct was cloned into pBRE, pHLT, or pGST vectors containing either thrombin or TEV protease cleavage sites to enable removal of the fusion tag after purification.

Since high solubility typically reflects correct folding and greatly simplifies downstream purification, we first assessed the expression behaviour of each fusion variant in *E. coli*. Analysis of total, soluble, and insoluble fractions by SDS-PAGE revealed that the His-tagged construct expressed strongly at the expected molecular weight (27.4 kDa) but was almost entirely insoluble (Figure 6.A), indicating accumulation in inclusion bodies. Introducing the HLT tag substantially improved solubility, shifting most of the protein to the soluble fraction at ∼36.5 kDa (Figure 6.B). The GST fusion tag also enhanced solubility (Figure 6.C), consistent with GST’s well-established role as a highly soluble carrier protein. These observations indicated that a solubilisation tag (HLT or GST) was required for successful production of the HasR dual-domain construct. The cleavable GST fusion protein confers another advantage besides solubilisation: it can be used as a separation step via affinity chromatography with glutathione resin, allowing for a “double-tag” (GST-HasR-His) purification strategy.

**Figure 6.**
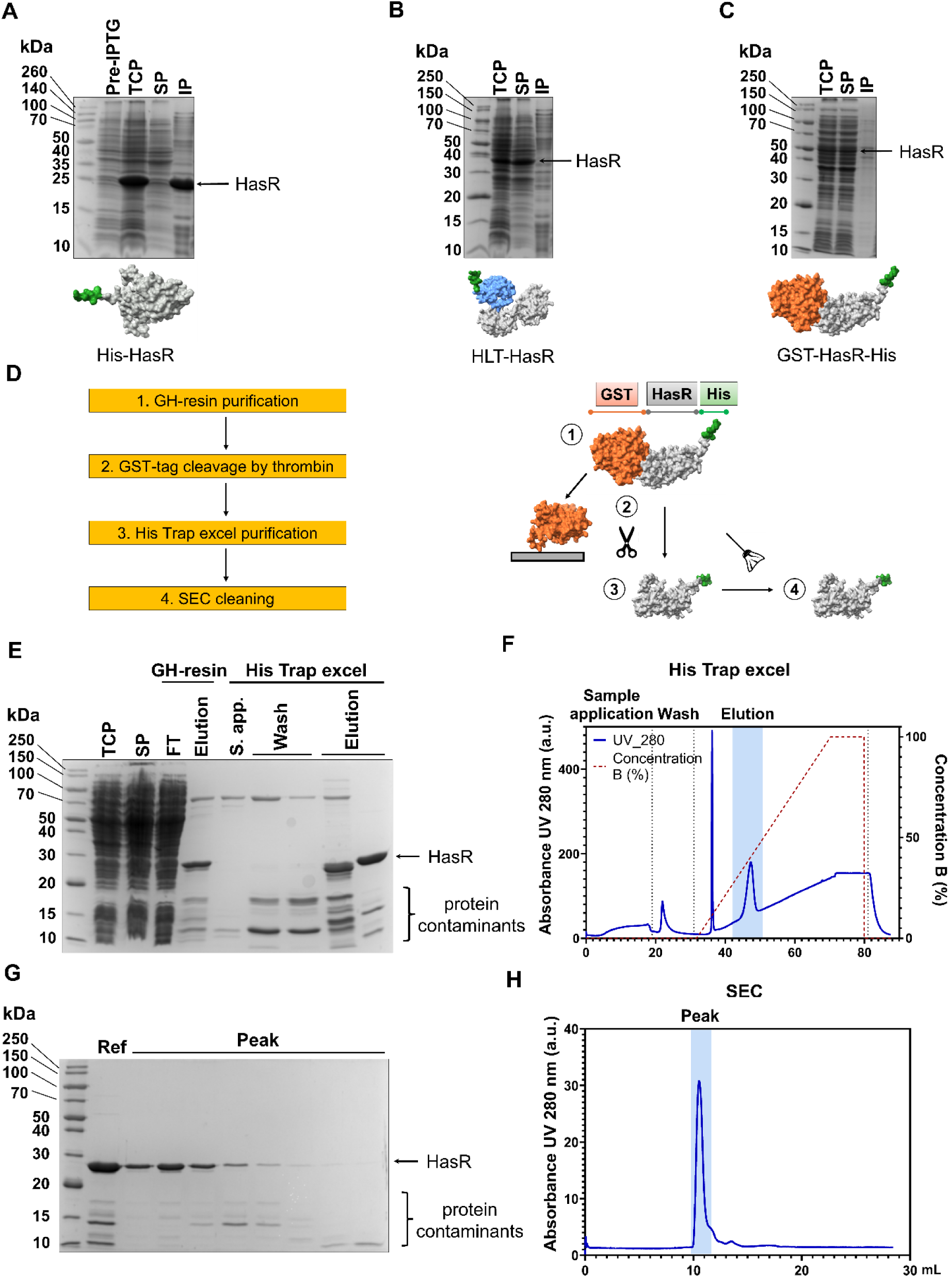
Experimental trials of the HasR construct expression with different fusion tags: **(A)** N-terminal 6-Histidine**, (B)** N-terminal HLT and **(C)** N-terminal GST with C-terminal 6-Histidine. HasR construct is denoted by the arrow [TCP= total cell protein lysate, SP= soluble protein fraction, IP= insoluble protein fraction]. HasR-His is primarily expressed in the insoluble fraction **(A)**, while the addition of the HLT tag **(B)** or GST **(C)** drives the expression to the soluble part. GST-HasR-His purification and characterization. **(D)** Flow chart of the GST/His purification process. The protein is first purified by means of GH-resin, where the GST-tag is cleaved by Thrombin, and then by affinity chromatography using His Trap excel column, as reported by **(E)** the SDS-PAGE and **(F)** the chromatogram from the AKTA. HasR construct band is denoted by arrow between 30 kDa and 20 kDa, some protein contaminants are present at lower molecular weight. The chromatogram reports Sample application, Wash and Elution phases; elution peak is highlighted by blue shadow. The protein is further purified by SEC, **(G)** SDS-PAGE and **(H)** the chromatogram from the AKTA are shown. HasR construct band is denoted by arrow between 30 kDa and 20 kDa, and a progressive reduction in protein contaminants is clearly visible. The chromatogram reports the elution phase, and the elution peak (Peak) is highlighted by blue shadow.

We next evaluated purification performance. The HLT-HasR construct was purified by nickel affinity chromatography. HLT-HasR expressed very well, resulting in a good protein yield, as indicated by the intense protein bands at the elution phase of the SDS-PAGE gel that represents approximately 70% of the total protein (Supplementary Figure S5.A). Nevertheless, the affinity purification step did not sufficiently purify the protein, as multiple additional proteins from the bacterial lysate were still present in the eluate (Supplementary Figure S5.A). Thrombin cleavage of the HLT tag released full-length HasR but also generated multiple degradation products (10–15 kDa), and prolonged digestion resulted in complete loss of intact protein (Supplementary Figure S5.B). This pronounced susceptibility to thrombin suggested non-specific proteolysis and made the HLT strategy unsuitable for further development.

Given these limitations, we focused on the dual-tag GST-HasR-His construct, which offered both enhanced solubility and the opportunity for sequential affinity purification. In addition, we evaluated TEV protease as an alternative cleavage strategy, anticipating greater specificity and reduced degradation compared with thrombin^29^.

### Recombinant HasR construct can be efficiently purified via a GST-tag and His-tag Affinity chromatography strategy

To purify the dual-domain GST-HasR-His construct, we established a sequential workflow designed to exploit both affinity tags and minimise contaminants. Cultures expressing the construct were grown and induced under conditions favouring soluble expression ^30^, and the soluble lysate was subjected to a three-step purification strategy (Figure 6.D).

In the first purification step (***Capture – Affinity Separation***), the GST tag enabled capture of the fusion protein on glutathione resin. The total cell lysate was incubated with glutathione resin, the unbound fractions were washed, and the GST fusion tag was cleaved with thrombin, releasing the HasR-His construct, while the GST remained bound to the resin. This step produced a clear HasR band with only minor additional species (Figure 6E).

The eluate was then subjected to a second, nickel affinity chromatography purification step (***Intermediate Purification – Affinity Chromatography***), which removed most remaining contaminants and yielded a dominant HasR band with a small number of lower-molecular-weight species attributable to trace impurities or partial degradation (Figure 6.E–F). The purity of HasR at this purification step was 79%.

The third, SEC purification step (***Polishing – Gel Filtration***) (Figure 6.G-H) removed residual contaminants and produced a single major peak corresponding to the HasR construct. ESI mass spectrometry confirmed the correct molecular weight and high purity (Supplementary Figure S2.C).

These results demonstrate that the dual-affinity GST-HasR-His strategy is markedly superior to the single-tag HLT-HasR approach, providing higher purity and fewer contaminants. It is a relatively quick and efficient method of purifying proteins of interest to satisfactory purity and can be followed with a final SEC polishing purification step to further elevate the purity. A key optimisation step involved the choice of protease for tag removal. Thrombin cleavage proved unsuitable, as the HasR construct was highly susceptible to non-specific digestion, resulting in extensive degradation and reduced yield (Supplementary Figure S5B). In contrast, TEV protease provided specific and efficient cleavage. On-resin TEV digestion followed by nickel affinity purification (Supplementary Figure S6A) increased purity to 87% and substantially reduced the presence of 10–20 kDa degradation products (Supplementary Figure S6B–C). The final yield of purified HasR construct was ∼0.5 mg/L (4.19 mg from 9 L culture).

### HasR construct adopts a partially fully folded conformation

Having purified the HasR dual-domain construct to high purity, we next assessed whether it adopted the expected folded architecture. Circular dichroism (CD) spectroscopy revealed a mixed secondary-structure profile comprising 20.3% α-helix, 27.0% right-twisted antiparallel β-strand, and 52.7% disordered content (Figure 7.A–B). These values closely match AlphaFold2 predictions for the combined domains (25% α-helix, 25% β-sheet, ∼50% disordered), indicating that the construct contains both structured and intrinsically flexible regions.

**Figure 7.**
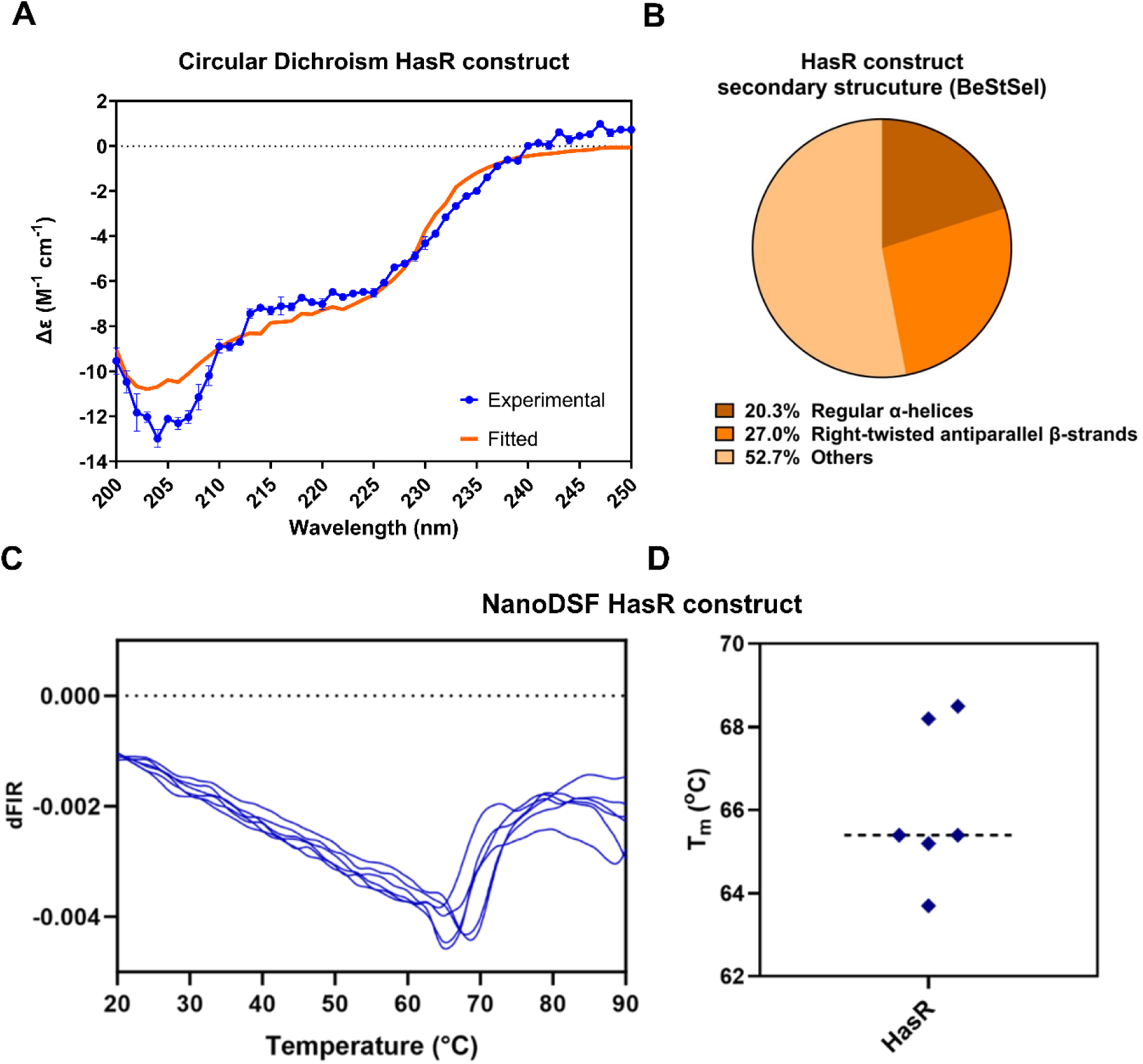
**(A)** Circular Dichroism spectrum of HasR construct measured in triplicate at 25°C between 200 and 250 nm (in blue), the fitted curve is also reported (in orange). **(B)** Secondary structure prediction by BeStSel server using the CD values, indicating a composition of 20.3% of regular α-helix (brown), 27.0% of right-twisted antiparallel β-strand (orange) and 52.7% others (disordered, pink) of HasR-His. **(C)** Unfolding of purified HasR construct determined by Nano Differential Scanning Fluorimetry (NanoDSF). The data shown are the first derivative of the ratio of fluorescence intensity read at 350 nm over that at 330 nm (dFIR (350 nm/330 nm)). The global minimum corresponds to the melting temperature of the protein, T_m_. **(D)** Extracted T_m_ values from the NanoDSF traces indicate that the protein is folded and that it has moderate thermal stability (66 ± 1.7°C), typical for many soluble, functional proteins. The NanoDSF experiments were performed with a Prometheus NanoDSF instrument (NanoTemper Technologies). Thermal denaturation was performed from 20°C to 90°C with a 1°C/min rate.

Thermal stability was evaluated using nanoDSF (Figure 7.C–D). Monitoring intrinsic tryptophan and tyrosine fluorescence during heating from 20°C to 90°C yielded a melting temperature of 66 ± 1.7°C. This Tm falls within the range typical for soluble bacterial proteins (53–73°C), suggesting that the folded regions of the construct are moderately stable ^31^.

To probe tertiary structure, we collected 1D NMR spectra of the purified protein (Figure 8.A). The spectrum displayed well-dispersed resonances characteristic of folded domains, alongside numerous amide signals in the random-coil region, consistent with the substantial disorder observed by CD. Together, these data indicate that the HasR construct contains a folded core with additional flexible regions.

**Figure 8.**
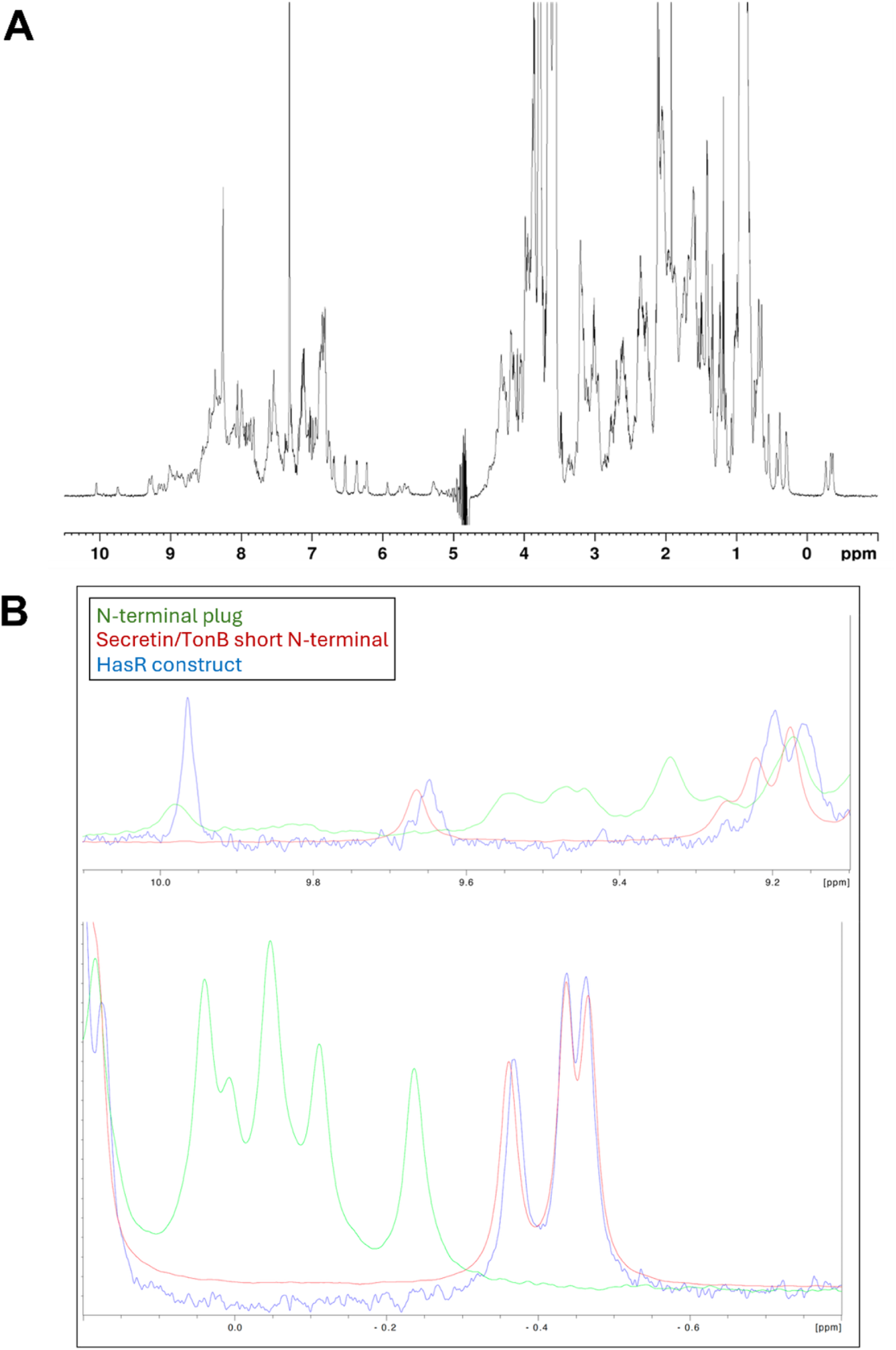
NMR spectra of HasR constructs and comparison with NMR spectra of the individual domains. **(A)** 1D ^1^H spectrum of the HasR construct on the 800 MhZ. **(B)** Superimposition of the spectra of the three constructs (N-terminal plug in green, Secretin/TonB short N-terminal in red and HasR construct in blue).

To determine whether both domains retained their native folds within the fusion construct, we compared NMR spectra of the individual domains with that of the combined protein (Figure 8.B). The resonances corresponding to the fully folded N-terminal plug domain (green trace) are not present in the spectrum of the HasR construct (blue trace), indicating the plug domain does not adopt the same well-packed β-sheet conformation when expressed together with the Secretin/TonB short N-terminal domain. In contrast, the resonances corresponding to the Secretin/TonB domain (red trace) were preserved, indicating that this domain remains stably folded in both contexts.

Finally, to obtain residue-level insight into structural stability, we performed Hydrogen/Deuterium Exchange Mass Spectrometry (HDX-MS). The protein was digested efficiently, yielding a coverage map of 88 peptides with 100% coverage and 5.02 average redundancy, as well as areas of very high redundancy (>10 peptides) (Supplementary Figure S7). H/D labelling across five timepoints spanning milliseconds to minutes (300 ms, 500 ms, 1 s, 30 s, 5 min) allows for capturing fast and slower exchange kinetics (Supplementary Figure S8). The exchange data for each peptide were averaged and represented as a heat map across the protein sequence, where yellow colour corresponds to lower deuterium uptake (stable/folded structure), and red colour corresponds to higher deuterium uptake (dynamic/unfolded structure).

HDX showed that the Secretin/TonB short N-terminal domain is a fully folded and stable region, as the deuterium labelling was low (<40%, yellow), indicating natively folded structure (Figure 9.A). The N-terminal plug domain showed increased deuterium uptake, reaching 62%, which corresponds well with the loop regions (amino acids 160-180) that are expected to be very flexible. Importantly, intermediate deuterium uptake was observed in the expected β-strand regions (amino acids 130-150, 200-220), indicating weakly stable structure, rather than complete disorder. Mapping HDX values onto the AlphaFold3 model of the construct (Figure 9B) and comparing them with annotated domain boundaries (Figure 9C) confirmed this domain-specific behaviour.

**Figure 9.**
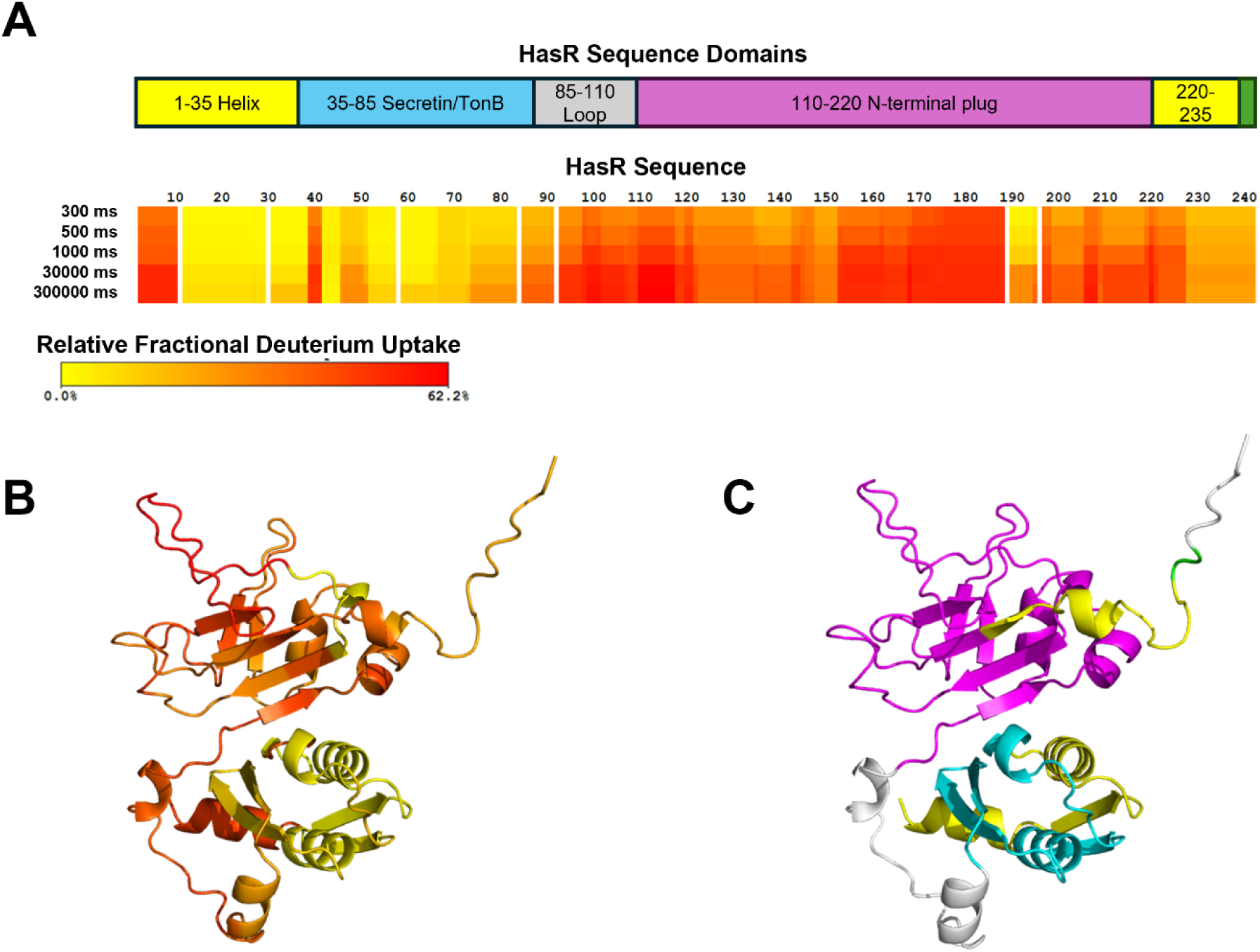
HDX-MS of HasR construct indicates folded structure. **(A)** Deuterium Uptake Heatmap across the sequence of the HasR construct. Areas in yellow indicate lower deuterium uptake, corresponding to folded and stabler areas of the protein. Areas in red indicate higher deuterium uptake, corresponding to more dynamic or unfolded areas of the sequence (e.g. loops). The protein domains (Helix, Secretin/TonB, N-terminal plug) are also highlighted. **(B)** Superimposition of the deuterium uptake data on the HasR construct 3D structure. **(C)** Highlighted sequence domains on the HasR construct 3D structure (corresponding to A), to aid comparison. Construct structure generated via AlphaFold.02.

Overall, the biophysical data demonstrate that the Secretin/TonB short N-terminal domain remains stably folded within the fusion construct, whereas the N-terminal plug domain becomes only weakly structured, with extensive flexibility in loop regions. These findings align with the NMR and CD results and reveal that domain context strongly influences the folding behaviour of HasR’s soluble regions.

## Discussion

In this study, we established a robust workflow for expressing, purifying, and characterising purifying soluble domains from membrane-associated bacterial receptors, using the soluble N-terminal domains of the TonB-dependent heme receptor HasR from *Pseudomonas aeruginosa* as a model system. By combining systematic tag screening, dual-affinity purification, and orthogonal biophysical analyses, we generated high-quality constructs suitable for downstream structural and ligand-discovery applications. Our findings highlight both the opportunities and the challenges inherent in working with isolated domains of membrane-embedded receptors.

A central outcome of our work is the identification of an effective solubilisation and purification strategy for the HasR dual-domain construct. Initial trials showed that simple His-tag fusion was insufficient, yielding predominantly insoluble protein. Introducing engineered solubility tags significantly improved expression behaviour, with both HLT and GST fusions driving the construct into the soluble fraction. Among these, the GST-HasR-His dual-tag design proved most advantageous, providing high solubility, compatibility with sequential affinity purification, and robust performance across multiple purification steps.

The dual-affinity approach (GST capture followed by His-tag enrichment) offered several methodological benefits. First, it enabled efficient removal of host proteins early in the process, reducing the burden on downstream chromatographic steps. Second, it allowed purification under mild, native conditions without the need for detergents, which is particularly valuable for membrane-associated domains that may be sensitive to denaturation. Third, the workflow was readily adaptable, permitting optimisation of protease cleavage conditions and integration of SEC polishing to achieve high purity.

The final purification strategy—GST affinity capture, TEV cleavage, His-tag enrichment, and SEC polishing—yielded high-purity protein suitable for downstream biophysical analysis. Importantly, this workflow is modular and broadly applicable. The combination of solubilisation tags, dual-affinity steps, and orthogonal polishing can be readily adapted to other membrane-associated receptors or challenging bacterial proteins. The approach is also compatible with large-scale production, as demonstrated by the reproducible yields obtained across multiple litres of culture.

Beyond purification, the study establishes a methodological framework for validating the structural integrity of soluble receptor domains. CD spectroscopy, nanoDSF, NMR, and HDX-MS together provide a comprehensive toolkit for assessing folding, stability, and suitability for downstream applications such as ligand screening. The integration of these techniques ensures that purified constructs are not only homogeneous but also structurally competent for functional studies. Importantly, the folding behaviour of the two HasR domains was consistent with expectations from individual expression trials: the Secretin/TonB short N-terminal domain remained well folded both as an isolated protein and within the dual-domain construct, demonstrating that its structural integrity is preserved across expression formats. In contrast, the N-terminal plug domain displayed conformation-dependent behaviour, adopting a stable fold when expressed alone but exhibiting weaker folding within the combined construct. These observations suggest that flexible loop regions within the plug domain exert a greater influence on its folding when the domain is expressed in the context of the larger construct. Together, these results highlight the value of incorporating orthogonal biophysical methods into purification workflows to ensure that recombinant receptor domains retain the structural features required for downstream biochemical and ligand-discovery studies.

## Conclusion

Overall, this study establishes a practical and adaptable methodology for producing soluble domains of membrane-embedded receptors. The combination of solubilisation tags, dual-affinity purification, specific protease cleavage, and comprehensive folding validation provides a reliable workflow that can be readily applied to other TonB-dependent receptors or challenging bacterial proteins. This approach enables access to high-quality protein suitable for biophysical and biochemical studies, facilitating future efforts in structural characterisation and therapeutic ligand discovery targeting bacterial iron-acquisition pathways.

## Materials and Methods

### Cloning of individual domains

The genes encoding for *P. aeruginosa* Secretin/TonB short N-terminal and N-terminal plug domains were obtained from the Pseudomonas Genome Database^35^. The gene sequences were designed with the *BamHI* restriction site at the 5ʹ position and with the 6-Histidine tag and the *HindIII* restriction site at the 3ʹ position. The genes were synthesized from IDT and codon optimized through GeneArt (Thermofisher Scientific) for *E. coli* cells. Both genes were cloned into pBRE vector harboring a 6-Histidine tag, ligation products were transformed into chemically competent Bronze *E. coli* cells via heat-shock treatment, and the positive clones were selected on 2xYT (Yeast Extract Tryptone) plates containing 100 µg/ml ampicillin. The success of the ligations was tested by enzymatic digestion with *BamHI* and *HindIII* and gel electrophoresis with 1% agarose gel in 1x TAE buffer. The accuracy of the constructs was confirmed by Sanger sequencing from GENEWIZ with primers for T7 promoter and T7 terminator. The alignment of HasR gene sequence with T7 promoter and T7 terminator reads was performed using SnapGene software.

### Expression and Purification of individual domains

Chemically competent Lucigen C41(DE3) *E. coli* cells were transformed with ligation products obtained from pBRE vector and Secretin/TonB short N-terminal and N-terminal plug domains. The transformation mixes were used to inoculate 10 ml 2xYT containing 100 µg/ml ampicillin, then grown overnight at 37°C and 220 rpm. The overnight cultures were used to inoculate the mains cultures with a 1:100 ratio and grown at 37°C and 220 rpm until an OD_600_ of 0.8 was measured. Protein expression was induced by the addition of isopropyl-b-D-1-thiogalactopyranoside (IPTG) to a final concentration of 1 mM. The mains cultures were then grown overnight at 20°C and 220 rpm. Bacterial pellets from main cultures were recovered by centrifugation (10000 g, 5 min, 4°C), resuspended in Buffer A (50 mM NaPO_4_ pH 7.5, 150 mM NaCl) supplemented with cOmplete™ Protease Inhibitor Cocktail and DNAase I (1 µg/ml) and lysed by Emulsiflex. Bacterial lysates were centrifugated at 35000 g, 35 min, 4°C, and the supernatants were recovered for protein purification. N-terminal plug and Secretin/TonB short N-terminal domains were purified starting from affinity chromatography with PureCube 100 Compact Cartridge Ni-INDIGO 1 ml, with a flow rate of 1 ml/min using an AKTA Pure system. After sample application, the column was washed with 30 CV in Buffer A2 (50 mM NaPO_4_ pH 7.5, 500 mM NaCl) supplemented with 20 mM imidazole, and the elution was performed with 40 CV of a linear gradient of the imidazole of the Buffer B from 0% to 100% followed by 10 CV at 100% Buffer B (50 mM NaPO_4_ pH 7.5, 150 mM NaCl, 500 mM imidazole). Protein aliquots corresponding to the elution peak were recovered, assessed using 15% sodium dodecyl sulfate-polyacrylamide gel electrophoresis (SDS–PAGE), and dialyzed against 1 liter Buffer A at 4°C, overnight. After dialysis, SEC with Superdex^TM^ 75 Increase 10/300 GL column was used protein contaminants coming from the bacterial lysate. The column was equilibrated overnight in Buffer A with 1.2 CV at 0.8 ml/min flow. After sample application, the elution was performed in Buffer A with 1.2 CV at 0.5 ml/min flow. Each protein aliquot corresponding to the elution peak was assessed using 15% SDS–PAGE. N-terminal plug domain purification was completed by using Ion Exchange chromatography (IEX) with MonoQ column using a gradient of NaCl from 150 mM to 2 M to elute the protein of interest. Protein concentrations were determined by UV adsorption set at 280 nm and adjusted with the predicted molar extinction coefficients calculated by ProtParam tool at ExPASy web portal (ɛ= 12950 M^−1^ cm^−1^ and ɛ= 2980 M^−1^ cm^−1^, for N-terminal plug and Secretin/TonB short N-terminal, respectively).

### HasR construct cloning

The gene encoding for *P. aeruginosa* HasR_44-276_ (including Secretin/TonB short N-terminal and N-terminal plug domains) was obtained from the Pseudomonas Genome Database^35^. The gene sequence was designed with the *BamHI* restriction site at the 5ʹ position and with the 6-Histidine tag and the *HindIII* restriction site at the 3ʹ position (Supplementary Figure S4-A and -C). The HasR gene was synthesized from IDT and codon optimized through GeneArt (Thermofisher Scientific) for *E. coli* cells. The gene was cloned into pGST vectors harboring a GST tag sequence and either a thrombin-or a TEV-cleavage site (Supplementary Figure S4-A, -B and -C). Similarly, HasR_44-276_ gene was amplified from gDNA of *P. aeruginosa PAO1* with the forward primer (5ʹ-TATA<u>GGATCC</u>AGCCAGCAGCAGACCGC-3ʹ) bringing the *BamHI* restriction site, underlined in the sequence, and the reverse primer (5ʹ-TAAT<u>AAGCTT</u>TTAACCGACCTGCTTGCCCG-3ʹ) bringing the *HindIII* restriction site, underlined in the sequence. The PCR product was cloned both into pBRE vector harboring a 6-Histidine tag and into pHLT vector harboring a Histidine Lipoyl tag (HLT)^27^. All four ligation products were transformed into chemically competent Bronze *E. coli* cells via heat-shock treatment, and the positive clones were selected on 2xYT plates containing 100 µg/ml ampicillin. The success of the ligation and the accuracy of the constructs were assessed as described above at *Cloning of individual domains*.

### HasR construct expression test

Chemically competent Lucigen C41(DE3) *E. coli* cells were transformed with pGST-thrombin-HasR-His and pGST-TEV-HasR-His constructs via heat shock treatment to produce a GST tag cleavable GST-HasR-His protein construct (Supplementary Figure S4-A). Similarly, the same *E. coli* cells were transformed with pBRE-HasR and pHLT-HasR constructs via heat shock treatment to produce a 6-Histidine tag cleavable His-HasR and a HLT cleavable HLT-HasR constructs, respectively. The transformation mix was used to inoculate 10 ml 2xYT containing 100 µg/ml ampicillin, then grown overnight at 37°C and 220 rpm. The overnight culture was used to inoculate the main culture with a 1:100 ratio and grown at 37°C and 220 rpm until an OD_600_ of 0.8 was measured. Protein expression was induced by the addition of IPTG to a final concentration of 1 mM. The main culture was then grown overnight at 20°C and 220 rpm. Bacterial pellet from 1 ml of main culture was recovered by centrifugation (10000 rpm, 2 min), resuspended in 1 ml of BugBuster® MasterMix (Novagen) and incubated for 30 min at room temperature with shaking. A 20 µl aliquot was saved as total cell protein lysate (TCP). After centrifugation (13000 rpm, 10 min) the supernatant was recovered as soluble protein fraction (SP); while the pellet was washed in 1 ml of 1:10 diluted BugBuster® MasterMix, dissolved in 300 µl of Solubilization buffer (100 mM Tris-HCl pH 8.2, 8 M urea, 100 mM β-mercaptoethanol) and centrifugated at 13000 rpm, 5 min to isolate the insoluble protein fraction (IP). The protein expression was analyzed using 15% SDS–PAGE.

### Purification of the HLT-HasR construct and screening of thrombin cleavage

HLT-HasR construct was purified by affinity chromatography with HisTrap^TM^ HP 1 ml column with a flow rate of 1 ml/min using an AKTA Pure system. After sample application, the column was washed with 10 CV in Buffer A2, and the elution was performed with 10 CV of a linear gradient of the imidazole of the Buffer B from 0 to 100 % followed by 5 CV at 100% Buffer B. Protein aliquots corresponding to the elution peak were assessed using 15% SDS–PAGE. The screening of the thrombin cleavage was performed with 4 U of bovine Thrombin (1U/µl, 7.5 mg/ml) per mg of HLT-HasR at room temperature with shaking.

### HasR construct purification and GST-tag removal

HasR construct was expressed as described above at *HasR construct expression test*; after centrifugation (10000 g, 15 min, 4°C), the bacterial pellet was resuspended in Buffer A supplemented with cOmplete™ Protease Inhibitor Cocktail and DNAase I (1 µg/ml) and lysed by Emulsiflex. Bacterial lysate was centrifugated at 35000 g, 35 min, 4°C, and the supernatant was recovered for protein purification (Purification I and Purification II).

### Purification I

Purification I was performed as described in Figure 6-D. Around 4 g of bacterial pellet were used to isolate HasR construct. The supernatant was incubated with 1 ml (0.5 g) GH-resin (Cambridge Bioscience LTD) for 2 hours, at 4°C with shaking. After centrifugation (500 g, 10 min) GH-resin was washed with 5 CV three times in Buffer A2 and resuspended in 4 ml of Buffer A. GST tag removal was performed on GH-resin by bovine thrombin cleavage with 10 U (1 U/µl, 7.5 mg/ml) of the protease at room temperature for 2 hours with shaking. The digestion mix including the GH-resin was transferred to a gravity column and the cleaved HasR construct was eluted by adding 2.5 CV three times of Buffer A. The cleaved product was further purified by affinity chromatography with HisTrap^TM^ excel 1 ml column with a flow rate of 1 ml/min using an AKTA Pure system. After sample application, the column was washed with 20 CV in Buffer A2 and the elution was performed with 40 CV of a linear gradient of the imidazole of the Buffer B from 0% to 100% followed by 10 CV at 100% Buffer B. Protein aliquots corresponding to the elution peak were recovered, assessed using 15% SDS–PAGE, and dialyzed against 1 liter Buffer A at 4°C, overnight. HasR construct was separated from digestion byproducts by SEC (Size Exclusion Chromatography). Superdex^TM^ 75 Increase 10/300 GL column was equilibrated overnight in Buffer A with 1.2 CV at 0.8 ml/min flow. After sample application, the elution was performed in Buffer A with 1.2 CV at 0.5 ml/min flow. Each protein aliquot corresponding to the elution peak was assessed using 15% SDS–PAGE. Protein concentration was determined by UV adsorption set at 280 nm and adjusted with the predicted molar extinction coefficient calculated by ProtParam tool at ExPASy web portal (ɛ= 15930 M^−1^ cm^−1^). Protein purity was calculated from SDS-PAGE gels, using the GelGenie plugin in QuPath, using the Universal model for segmentation, as previously described^36^.

### ESI-MS

Electrospray ionisation mass spectrometry was performed on a Xevo G2 mass spectrometer with data analysed using MassLynx software (Waters UK) (Yusuf Hamied Department of Chemistry, University of Cambridge, UK).

### Purification II

Purification II was performed as described in Supplementary Figure S6-A. Around 140 g of bacterial pellet were used to isolate HasR construct. The supernatant was incubated with 6×1 ml GH-resins (0.5 g) for 2 hours, at 4°C with shaking. After centrifugation (500 g, 10 min) each GH-resin was washed with 5 CV three times in Buffer A2 and resuspended in 2 ml of Buffer A. GST tag removal was performed on GH-resin by biotin-TEV cleavage with 240 U (10 U/µl) of the protease at 4°C, overnight with shaking. The digestion mix including the GH-resin was transferred to a gravity column and the cleaved HasR construct was eluted by adding 2.5 CV three times of Buffer A. GST tag was eluted from GH-resin by adding 5 CV three times of Elution Buffer A (50 mM NaPO_4_ pH 8, 150 mM NaCl, 20 mM reduced glutathione). The cleaved product was further purified by affinity chromatography with PureCube 100 Compact Cartridge Ni-INDIGO 1 ml with a flow rate of 1 ml/min using an AKTA Pure system. After sample application, the column was washed with 30 CV in Buffer A2 supplemented with 20 mM imidazole, and the elution was performed with 40 CV of a linear gradient of the imidazole of the Buffer B from 0% to 100% followed by 10 CV at 100% Buffer B. Protein aliquots corresponding to the elution peak were recovered, assessed using 15% SDS–PAGE, and dialyzed against 1 liter Buffer A at 4°C, overnight. Protein concentration was determined by UV adsorption set at 280 nm and adjusted with the predicted molar extinction coefficient calculated by ProtParam tool at ExPASy web portal (ɛ= 15930 M^−1^ cm^−1^).

### Circular Dichroism (CD)

CD was measured on Chirascan CD spectrometer (Applied Photophysics, Leatherhead, UK) in 1-mm-pathlength Precision Cells (110-QS;), using 6 µM of HasR construct, 11 µM of N-terminal plug and 17 µM of Secretin/TonB short N-terminal solutions in Buffer A in a 10 mm path length cuvette (Hellma Analytics, Müllheim, Germany). CD spectrum was registered three times in the range of 200-250 nm at 25°C. Beta Structure Selection (BeStSel) server was used to determine the protein secondary structure and fold recognition from CD spectrum, giving an indication of α-helix and β-sheets content^37^.

### NMR (1D spectrum)

The NMR samples (HasR construct concentration = 155 µM, N-terminal plug domain concentration = 86 µM and Secretin/TonB domain concentration = 352 µM) were prepared in phosphate buffer (20 mM KPi (Potassium Phosphate) + 150 mM NaCl) with 5% D_2_0. 1D ^1^H spectra were recorded at 25°C, with water suppression using excitation sculpting with gradients^38^ on a Bruker Avance III HD 800 MHz spectrometer.

### Nano Differential Scanning Fluorimetry (NanoDSF)

NanoDSF was performed with a Prometheus NanoDSF instrument (NanoTemper Technologies) to evaluate the melting temperature of the HasR construct. Protein thermal stability was estimated by calculating the first derivative of the ratio of fluorescence intensity read at 350 nm over 330 nm (dFIR (350 nm/330 nm)) from 20°C to 90°C with a 1°C/min rate.

### Hydrogen–deuterium exchange Mass Spectrometry (HDX-MS)

Hydrogen–deuterium exchange (HDX) is performed using a fully automated, millisecond HDX labeling and online quench-flow instrument, ms2-min (Nicoya Lifesciences, CA), connected to an HDX manager (Waters). Samples in the equilibrium buffer are delivered into the labeling mixer and diluted 20-fold with labeling buffer at 20°C, initiating HDX. The duration of the HDX labeling depends on the mixing loops of varying length in the sample chamber of the ms2 min and the velocity of the carrier buffer, calibrated to a precision of 1 ms. The protein is labeled for a range of times from 300 ms to 5 min. Immediately postlabeling, the labeled sample is mixed with quench buffer in a 1:1 ratio at 0 °C in the quench mixer to arrest HDX. The sample is then centered on the HPLC injection loop of the ms2 min and sent to the HDX manager. Protein samples are digested onto an enzymate immobilized pepsin column (Waters) to form peptides. The peptides are trapped on a VanGuard 2.1 mm × 5 mm ACQUITY BEH C18 column (Waters) for 3 min at 175 μL/min and separated on a 1 mm × 100 mm ACQUITY BEH 1.7 μm C18 column (Waters) with a 7 min linear gradient of acetonitrile (5–40%) supplemented with 0.1% formic acid. The eluted peptides are analyzed on a Synapt G2-Si mass spectrometer (Waters). Deuterium incorporation into the peptides is measured in DynamX 3.0 (Waters) for peptides identified by separate HDMS^E^ experiments analysed in ProteinLynx Global Server 3.03 (Waters).

## Acknowledgments

I.M. acknowledges funding from the Royal Society (URF/R1/221795, IES∖R3∖223128, IES∖R2∖222107, RGS∖R1∖231266), the National Biofilms Innovation Centre (BB/R012415/1 03PoC20-105) and the David James Trust.

M.Z. acknowledges funding from Oppenheimer Fellowship (School of the Physical Sciences, University of Cambridge).

L.S.I. and P.J.E.R. acknowledge the support of Human Frontiers Science Program grant RGP0027/2020.

J.J.P. and M.K. acknowledge funding from a UKRI Future Leaders Fellowship (MR/Z000157/1, MR/T02223X/1).

This work was supported by the Francis Crick Institute through provision of access to the MRC Biomedical NMR Centre. The Francis Crick Institute receives its core funding from Cancer Research UK (CC1078), the UK Medical Research Council (CC1078), and the Wellcome Trust (CC1078).

