## Supplementary Information for "A generalisable method for the purification and biophysical characterisation of bacterial membrane receptors"

### Supplementary Materials

### A

#### **Aminoacidic sequence of HasR** (891 aa)

Signal peptide sequence (1-36 aa)

Secretin/TonB, short N-terminal domain (75-126 aa)

N-terminal plug domain (146-272 aa)

β-barrel domain and loops (273-891 aa)

Heme-coordinating extracellular loop (605-647 aa)

MKHRGWSAVRGGRKGAQLALGLGLVLLGTAALPLHAQDGADSASQQQTALRRVRLDIPA
QPLNRALLRFAEQAGVQVFFDSQRFAGLGSAAVHGEYLLADGLSQMLQGSPVEYRFSGK
DQLSLIRVSQDDLVQMSPSVISAARPDDWVYQTPHSVSVIGREQIERNPPRHAADMLEET
PGVYSSVSQQDPGLSVNIRGIQDYGRVNMSVDGMRQNYQQSGHQQRNGTLYVDPELLS
EVVIDKGASSAMGGAGVIGGIANFRTLEARDLVRPGKQVGGRVRLTSGLGGDANGTHFIG
SAAFAIGTEVWDMLVAASERHLGDYDPGTKGSIGELRTGAWFNPEAGQRVKHSPVAYSGY
VMRSRLAKLGVALPQDQRLQFSYLTTQVSYDDANMLNTENQALWEKLGSSDVRAQNFAID
YGYAPDNPLVDFKAKLYYVDNRNRQQTLQRGITPGYSITYQTDTYGAQAQNTSTFALDDLS
TLRANYGLEFFYDKVRPDSSQPRASTSAVGFPAAEGMTPKGDRALGSLFARLDYDYDDW
LNLNAGLRYDRYRLRGDTGFNARTFILGTTRQTDMPLQYAVDREEGRFSPTFGLSVKPGV
DWLQLFATYGKGWRPPAVTESLITGRPHGGGAENMYPNPFLSPERSKAWEVGFNVLKEN
LWFSDDRLGLKVAYFDTRVDDFIFMGMGMQPPGYGMAGIGNSAYVNNLDSTRFRGVEYQ
LDYDAGLAYGQLSYTHMIGSNDFCSKTAWLGGVTQTVKGSGRRPPVIDMRPDEQANAAT
HCSAVLGSAEHMPMDRGSLTLGMRFFDRRLDVGARARYSEGYSVAGGATVSQAGVYPA
DWKEYTVYDLYGSYRVSDELTLRLAMENVTDRAYLVPLGDVLAFTLGRGRTLQGTLEYQF

### B

**HasR construct (underlined, 44-276 aa)**

Secretin/TonB, short N-terminal domain (75-126 aa)

N-terminal plug domain (146-272 aa)

```
        10         20         30         40         50         60
MKHRGWSAVR GGRKGAQLAL GLGLVLLGTA ALPLHAQDGA DSASQQQTAL RRVRLDIPAQ
        70         80         90        100        110        120
PLNRALLRFA EQAGVQVFFD SQRFAGLGSA AVHGEYLLAD GLSQMLQGSP VEYRFSGKDQ
       130        140        150        160        170        180
LSLIRVSQDD LVQMSPSVIS AARPDDWVYQ TPHSVSVIGR EQIERNPPRH AADMLEETPG
       190        200        210        220        230        240
VYSSVSQQDP GLSVNIRGIQ DYGRVNMSVD GMRQNYQQSG HQQRNGTLYV DPELLSEVVI
       250        260        270        280        290        300
DKGASSAMGG AGVIGGIANF RTLEARDLVR PGKQVGGRVR LTSGLGGDAN GTHFIGSAAF
       310        320        330        340        350        360
AIGTEVWDML VAASERHLGD YDPGTKGSIG ELRTGAWFNP EAGQRVKHSP VAYSGYVMRS
       370        380        390        400        410        420
RLAKLGVALP QDQRLQFSYL TTQVSYDDAN MLNTENQALW EKLGSSDVRA QNFAIDYGYA
       430        440        450        460        470        480
PDNPLVDFKA KLYYVDNRNR QQTLQRGITP GYSITYQTDT YGAQAQNTST FALDDLSTLR
       490        500        510        520        530        540
ANYGLEFFYD KVRPDSSQPR ASTSAVGFPA AEGMTPKGDR ALGSLFARLD YDYDDWLNLN
       550        560        570        580        590        600
AGLRYDRYRL RGDTGFNART FILGTTRQTD MPLQYAVDRE EGRFSPTFGL SVKPGVDWLQ
       610        620        630        640        650        660
LFATYGKGWR PPAVTESLIT GRPHGGGAEN MYPNPFLSPE RSKAWEVGFN VLKENLWFSD
       670        680        690        700        710        720
DRLGLKVAYF DTRVDDFIFM GMGMQPPGYG MAGIGNSAYV NNLDSTRFRG VEYQLDYDAG
       730        740        750        760        770        780
LAYGQLSYTH MIGSNDFCSK TAWLGGVTQT VKGSGRRPPV IDMRPDEQAN AATHCSAVLG
       790        800        810        820        830        840
SAEHMPMDRG SLTLGMRFFD RRLDVGARAR YSEGYSVAGG ATVSQAGVYP ADWKEYTVYD
       850        860        870        880        890
LYGSYRVSDE LTLRLAMENV TDRAYLVPLG DVLAFTLGRG RTLQGTLEYQ F
```

**HasR individual domains (underlined)**

Secretin/TonB, short N-terminal domain (44-145 aa)

N-terminal plug domain (146-272 aa)

```
        10         20         30         40         50         60
MKHRGWSAVR GGRKGAQLAL GLGLVLLGTA ALPLHAQDGA DSASQQQTAL RRVRLDIPAQ
        70         80         90        100        110        120
PLNRALLRFA EQAGVQVFFD SQRFAGLGSA AVHGEYLLAD GLSQMLQGSP VEYRFSGKDQ
       130        140        150        160        170        180
LSLIRVSQDD LVQMSPSVIS AARPDDWVYQ TPHSVSVIGR EQIERNPPRH AADMLEETPG
       190        200        210        220        230        240
VYSSVSQQDP GLSVNIRGIQ DYGRVNMSVD GMRQNYQQSG HQQRNGTLYV DPELLSEVVI
       250        260        270        280        290        300
DKGASSAMGG AGVIGGIANF RTLEARDLVR PGKQVGGRVR LTSGLGGDAN GTHFIGSAAF
       310        320        330        340        350        360
AIGTEVWDML VAASERHLGD YDPGTKGSIG ELRTGAWFNP EAGQRVKHSP VAYSGYVMRS
       370        380        390        400        410        420
RLAKLGVALP QDQRLQFSYL TTQVSYDDAN MLNTENQALW EKLGSSDVRA QNFAIDYGYA
       430        440        450        460        470        480
PDNPLVDFKA KLYYVDNRNR QQTLQRGITP GYSITYQTDT YGAQAQNTST FALDDLSTLR
       490        500        510        520        530        540
ANYGLEFFYD KVRPDSSQPR ASTSAVGFPA AEGMTPKGDR ALGSLFARLD YDYDDWLNLN
       550        560        570        580        590        600
AGLRYDRYRL RGDTGFNART FILGTTRQTD MPLQYAVDRE EGRFSPTFGL SVKPGVDWLQ
       610        620        630        640        650        660
LFATYGKGWR PPAVTESLIT GRPHGGGAEN MYPNPFLSPE RSKAWEVGFN VLKENLWFSD
       670        680        690        700        710        720
DRLGLKVAYF DTRVDDFIFM GMGMQPPGYG MAGIGNSAYV NNLDSTRFRG VEYQLDYDAG
       730        740        750        760        770        780
LAYGQLSYTH MIGSNDFCSK TAWLGGVTQT VKGSGRRPPV IDMRPDEQAN AATHCSAVLG
       790        800        810        820        830        840
SAEHMPMDRG SLTLGMRFFD RRLDVGARAR YSEGYSVAGG ATVSQAGVYP ADWKEYTVYD
       850        860        870        880        890
LYGSYRVSDE LTLRLAMENV TDRAYLVPLG DVLAFTLGRG RTLQGTLEYQ F
```

**Figure S1.** HasR aminoacidic sequence and domains. **(A)** Schematic of the aminoacidic sequence of HasR, highlighting the different domains: signal peptide (in grey), Secretin/Ton B short N-terminal (in cyan), N-terminal plug (in purple), β-barrel and loops (in dark green), and heme-coordinating extracellular loop (in light green). Heme binding residues are highlighted in yellow: H221, H624 and I694. **(B)** Aminoacidic sequences of HasR construct and HasR individual domains used in this work.

A

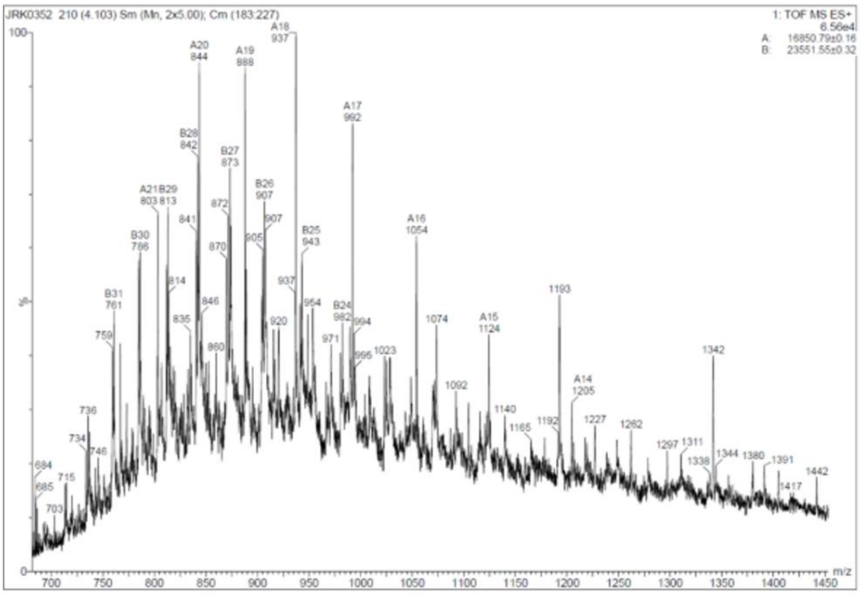

JRK0352 210 (4.103) Sm (Mn, 2x5.00); Cm (183:227)
1: TOF MS ES+
6.56e4
A: 16850.79±0.16
B: 23551.55±0.32
100
%
0
m/z
A20 844
A19 888
A18 937
A17 992
A16 1054
A15 1124
A14 1205
A21 803
B28 842
B27 873
B26 907
B25 943
B24 982
B29 813
B30 786
B31 761
1193
1342


B

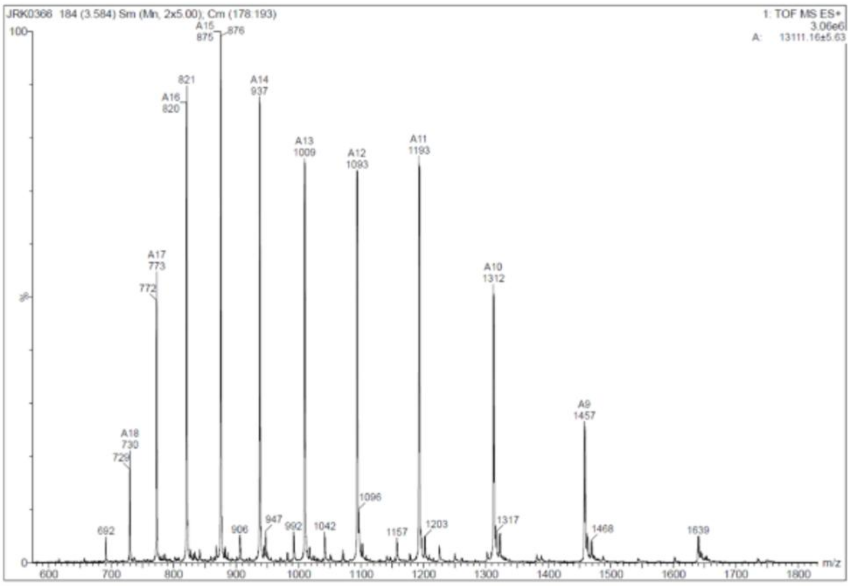

JRK0366 184 (3.584) Sm (Mn, 2x5.00); Cm (178:193)
1: TOF MS ES+
3.06e6
A: 13111.16±5.63
100
%
0
m/z
A15 875
A16 820
A14 937
A13 1009
A12 1093
A11 1193
A17 773
A10 1312
A9 1457
A18 730


C

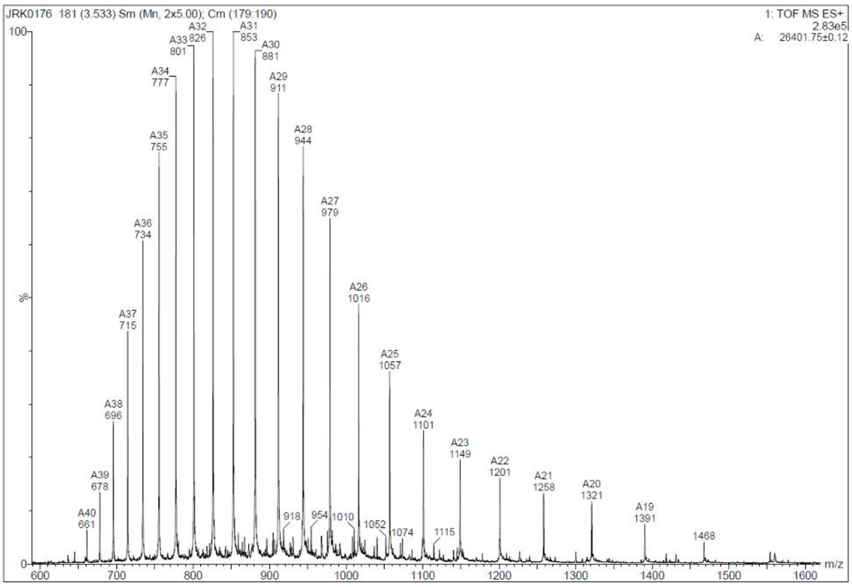

JRK0176 181 (3.533) Sm (Mn, 2x5.00); Cm (179:190)
1: TOF MS ES+
2.83e5
A: 26401.75±0.12
100
%
0
m/z
A32 826
A31 853
A33 801
A30 881
A34 777
A29 911
A35 755
A28 944
A27 979
A36 734
A26 1016
A37 715
A25 1057
A38 696
A24 1101
A23 1149
A22 1201
A39 678
A21 1258
A20 1321
A40 661
A19 1391

**Figure S2.** Electrospray ionisation mass spectrometry of N-terminal plug, Secretin/TonB short N-terminal domains and HasR construct. **(A)** The N-terminal plug domain showed a major peak at 16.9 kDa, close to the expected size of 15.8 kDa, and a minor peak at 23.6 kDa. **(B)** The Secretin/TonB short N-terminal domain size of 13.1 kDa was confirmed. **(C)** The HasR construct size of 26.4 kDa was also confirmed.

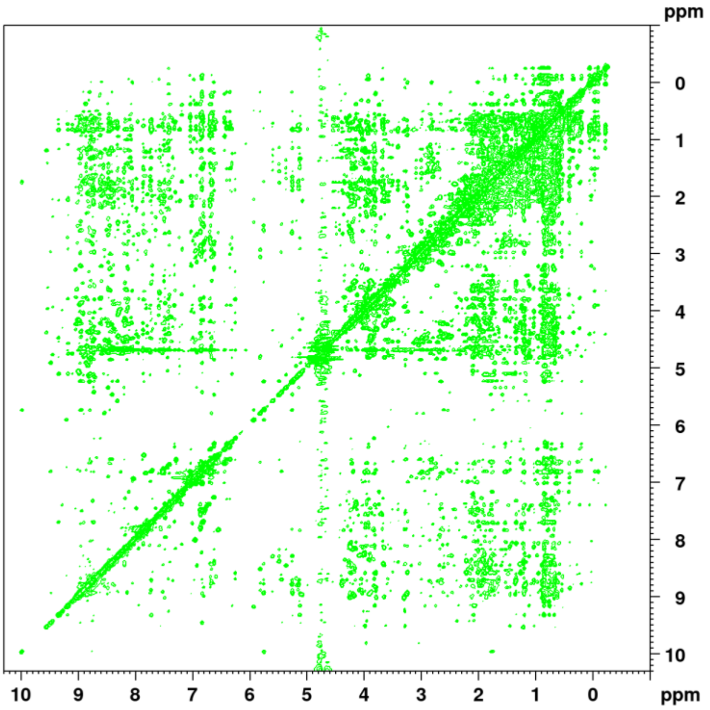


**Figure S3:** 2D NOESY NMR spectra of the N-terminal plug domain showing long-range, through-space contacts that are only present in a folded structure.

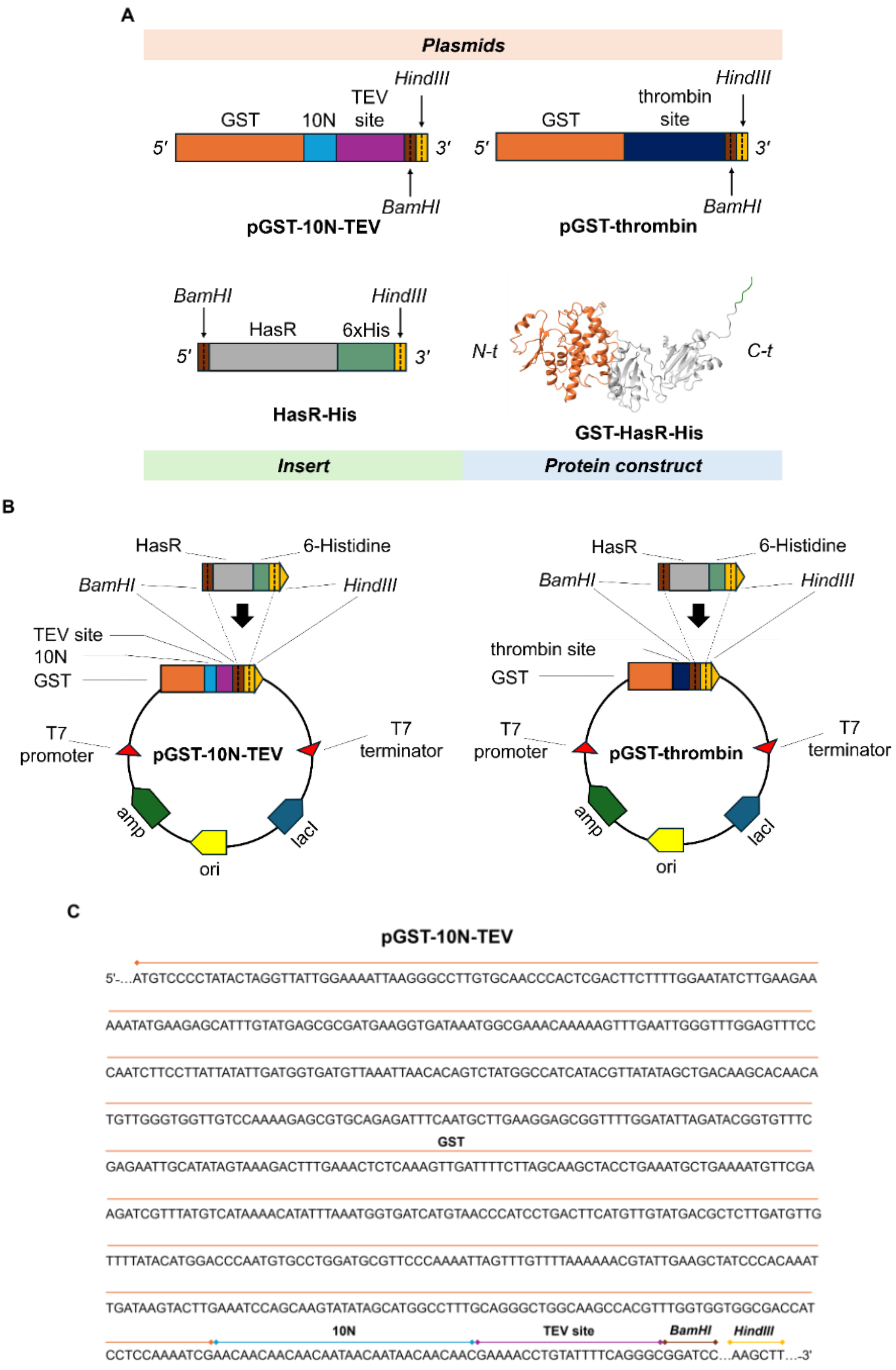
A
Plasmids
GST
10N
TEV
site
HindIII
5′
3′
BamHI
pGST-10N-TEV
GST
thrombin
site
HindIII
5′
3′
BamHI
pGST-thrombin
BamHI
HindIII
HasR
6xHis
5′
3′
HasR-His
N-t
C-t
GST-HasR-His
Insert
Protein construct
B
HasR
6-Histidine
BamHI
HindIII
TEV site
10N
GST
T7
promoter
pGST-10N-TEV
T7
terminator
amp
ori
lacI
HasR
6-Histidine
BamHI
HindIII
thrombin site
GST
T7
promoter
pGST-thrombin
T7
terminator
amp
ori
lacI
C
pGST-10N-TEV
5'-...ATGTCCCCTATACTAGGTTATTGGAAAATTAAGGGCCTTGTGCAACCCACTCGACTTCTTTTGGAATATCTTGAAGAA
AAATATGAAGAGCATTTGTATGAGCGCGATGAAGGTGATAAATGGCGAAACAAAAAGTTTGAATTGGGTTTGGAGTTTCC
CAATCTTCCTTATTATATTGATGGTGATGTTAAATTAACACAGTCTATGGCCATCATACGTTATATAGCTGACAAGCACAACA
TGTTGGGTGGTTGTCCAAAAGAGCGTGCAGAGATTTCAATGCTTGAAGGAGCGGTTTTGGATATTAGATACGGTGTTTC
GST
GAGAATTGCATATAGTAAAGACTTTGAAACTCTCAAAGTTGATTTTCTTAGCAAGCTACCTGAAATGCTGAAAATGTTCGA
AGATCGTTTATGTCATAAAACATATTTAAATGGTGATCATGTAACCCATCCTGACTTCATGTTGTATGACGCTCTTGATGTTG
TTTTATACATGGACCCAATGTGCCTGGATGCGTTCCCAAAATTAGTTTGTTTTAAAAAACGTATTGAAGCTATCCCACAAAT
TGATAAGTACTTGAAATCCAGCAAGTATATAGCATGGCCTTTGCAGGGCTGGCAAGCCACGTTTGGTGGTGGCGACCAT
10N
TEV site
BamHI
HindIII
CCTCCAAAATCGAACAACAACAACAATAACAATAACAACAACGAAAACCTGTATTTTCAGGGCGGATCC...AAGCTT...-3'

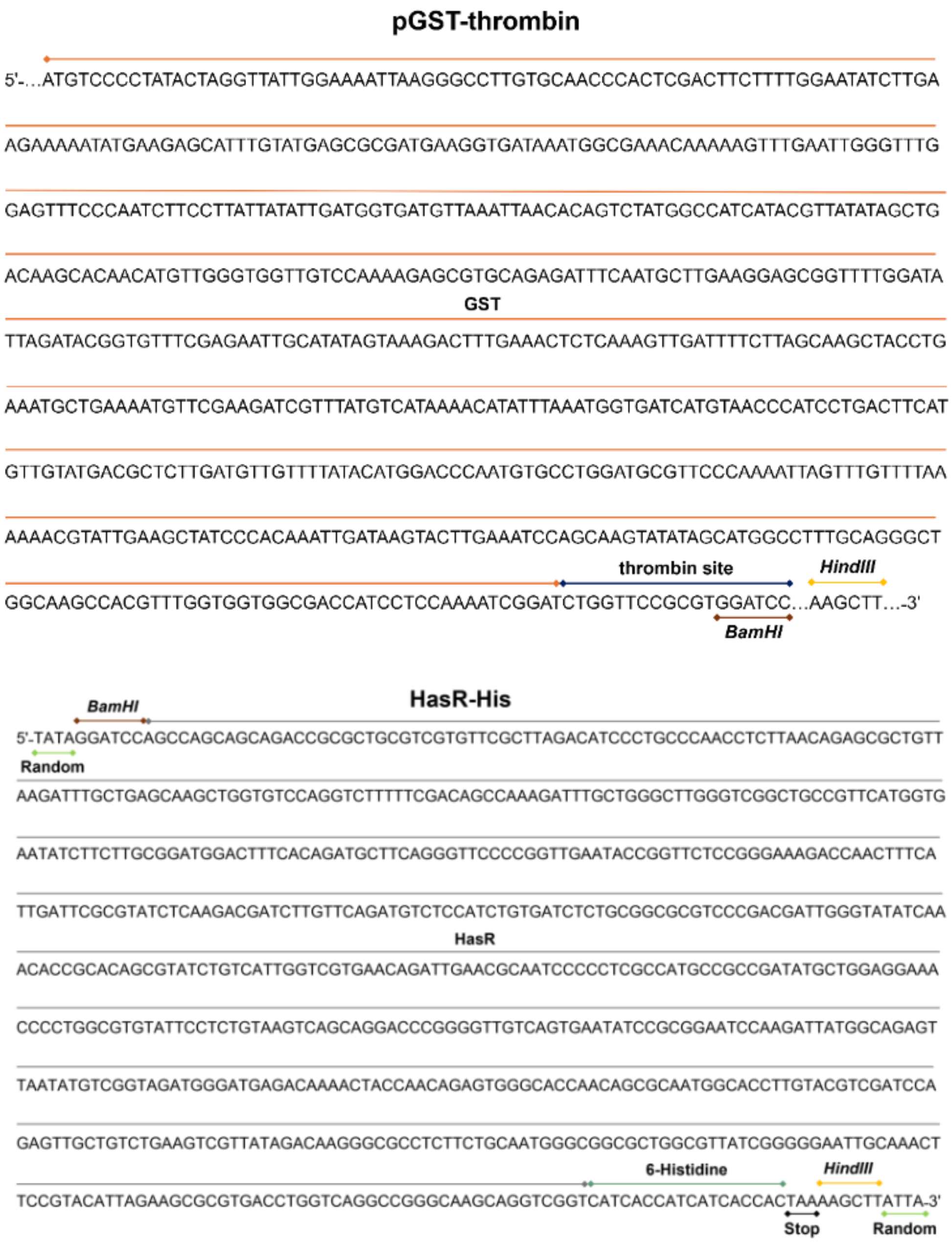


**Figure S4.** Illustration about the cloning of HasR-His construct in pGST vectors, which includes: **(A)** scheme of the plasmid vectors (pGST-10N-TEV and pGST-Thrombin), the insert (HasR-His) and the protein construct (GST-HasR-His). **(B)** Maps of the plasmid vectors (pGST-10N-TEV and pGST-Thrombin). **(C)** DNA sequences of plasmid vectors (pGST-10N-TEV and pGST-Thrombin) and the insert (HasR-His). HasR-His construct is cloned in the plasmid vectors by means of *BamHI* and *HindIII,* downstream of the GST, and of the TEV and thrombin cleavage sites.

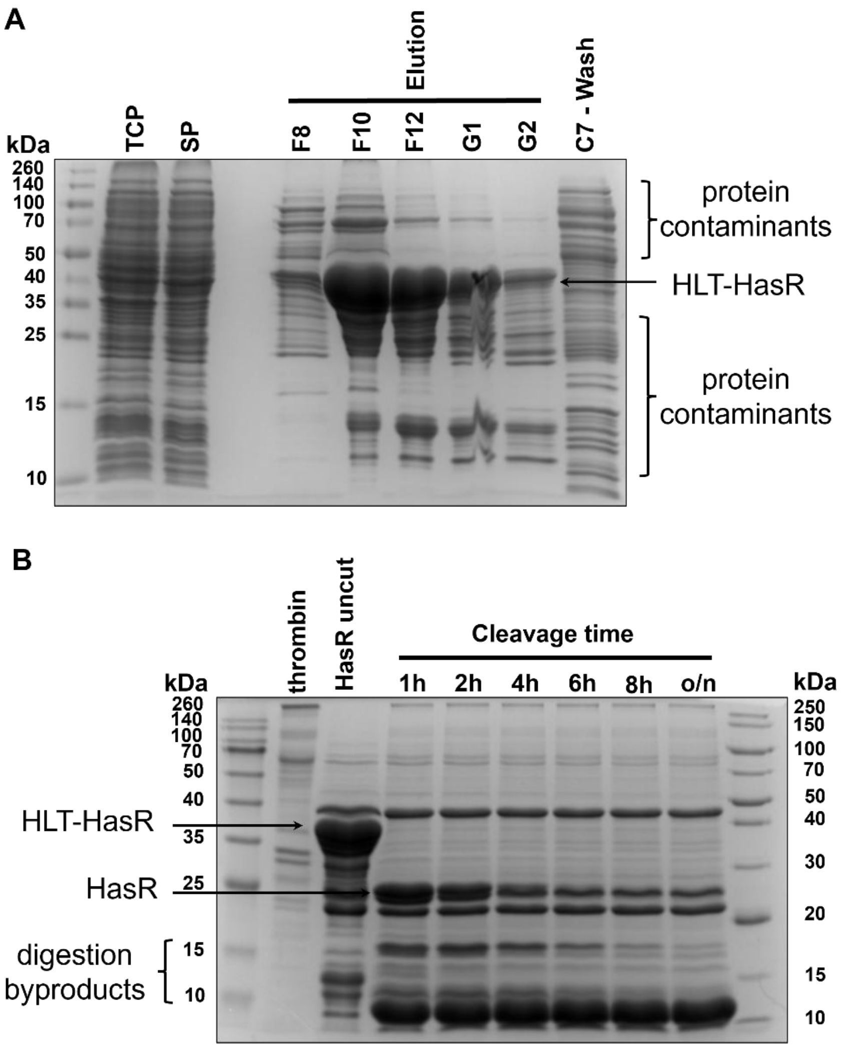


**Figure S5.** HLT-HasR construct purification and screening of the thrombin cleavage of the HLT linker. **(A)** Purification of HLT-HasR construct by affinity chromatography with HisTrap$^{TM}$ HP 1 ml column. Protein band is denoted by the arrow and protein contaminants at both higher and lower molecular weight are also indicated. **(B)** Screening of the thrombin cleavage of HLT-HasR construct at 1-2-4-6-8 hours and overnight, HLT-HasR and cleaved HasR protein bands are denoted by arrows, digestion byproducts are also reported.

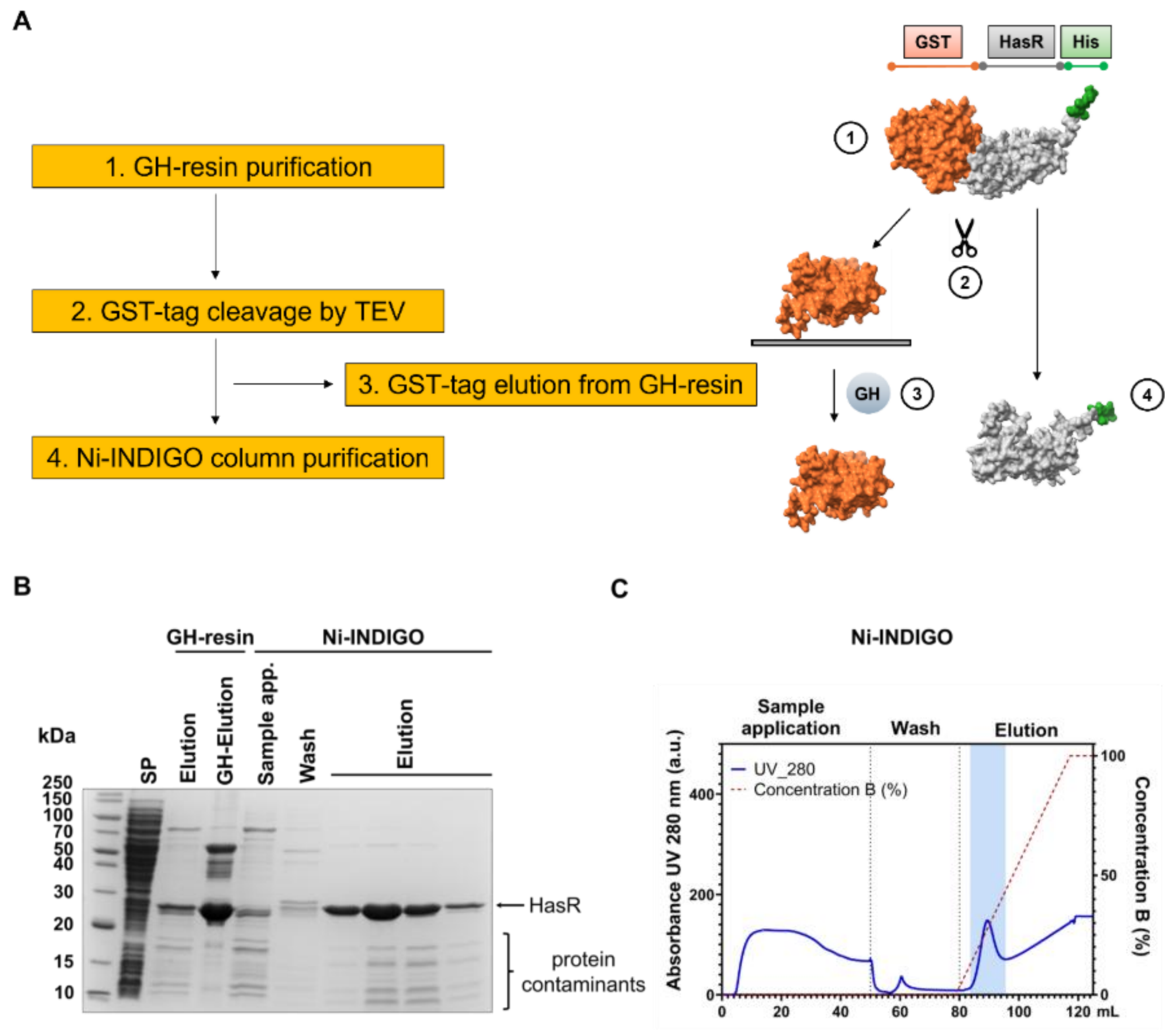


**Figure S6**. Purification II of GST-HasR-His construct. **(A)** Scheme about the purification process. **(B)** The protein is first purified by means of GH-resin, where the GST-tag is cleaved by TEV, and then **(B and C)** by affinity chromatography using Ni-INDIGO column, as reported by the SDS-PAGE and the chromatogram from the AKTA. HasR construct band is denoted by the arrow, protein contaminants are also indicated. [SP= soluble protein fraction, GH= glutathione].

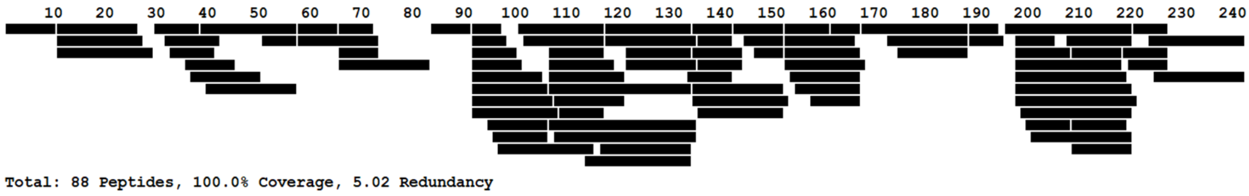


**Figure S7.** HDX-MS coverage map of HasR construct. Full sequence coverage (100%) and average 5.02 Redundancy (number of peptides an amino acid appears in) achieved.
